# The genomic outcomes of hybridization vary over time within a monkeyflower radiation

**DOI:** 10.1101/2024.08.26.609732

**Authors:** Aidan W Short, Matthew A Streisfeld

## Abstract

The accumulation of genetic differences through time can lead to reproductive isolation between populations and the origin of new species. However, hybridization between emerging species can occur at any point before isolation is complete. The evolutionary consequences of this hybridization may vary depending on when it occurred. If hybridization occurred later during the process, when ecological and genetic differences have accumulated between diverging lineages, low hybrid fitness can result in selection against gene flow. If hybridization occurred earlier, when barriers present were too weak to limit introgression, then hybridization can lead to genetic swamping. Alternatively, adaptive introgression can occur at any point during speciation. Thus, by understanding the history and genomic consequences of hybridization at different points along the speciation continuum, we can begin to understand how variation present within populations translates to divergence between species. Here, we identified the genomic signals of introgressive hybridization at different points during the divergence of two monkeyflower taxa endemic to the Channel Islands of California. We found that both ancient and recent introgression have shaped their genomes, but the impacts of selection on this foreign material varied. There was no signal of selection against ancient introgression, but we did find strong evidence for selection against recent introgression, potentially because there are more reproductive barriers in place now, reducing fitness in recent hybrids. Thus, this study reveals that hybridization can occur at multiple points throughout the divergence history of a radiation, but the processes shaping genome wide levels of introgression can change over time.

## 1 INTRODUCTION

Phenotypic and genetic differences that evolve between populations can lead to the accumulation of reproductive barriers and the origin of new species (Mayr 1942, Coyne & Orr 2004). Determining how reproductive isolation develops through time is critical for understanding how variation within populations translates to divergence between species (Matute & Cooper 2021). Although previously believed to be rare, hybridization between emerging species is now known to be common (Roux et al. 2016, Martin & Jiggins 2017, Malinsky et al. 2018, Stankowski et al. 2019, Liu et al. 2022). Depending on when it occurs, hybridization and subsequent gene flow between diverging lineages (i.e., introgression) can have various evolutionary consequences. For example, introgression can expose genetic incompatibilities that promote speciation (Coughlan & Matute 2020), it can swamp out divergence (Todesco et al. 2016), or in some cases, it can lead to the transfer of beneficial alleles across taxonomic boundaries (Suarez-Gonzalez et al. 2018). Thus, understanding the evolutionary history and genomic consequences of introgressive hybridization can help reveal the processes contributing to divergence and speciation (Harrison & Larson 2016, Ravinet et al. 2017).

In their classic study, Coyne & Orr (1989) found that reproductive isolation accumulated with levels of sequence divergence between multiple pairs of *Drosophila* species, a finding that has been supported in various other groups of divergent taxa (Moyle et al. 2004; Merot et al. 2017). Given that reproductive isolation takes time to arise, hybridization can occur at multiple points throughout the divergence of any pair of species (e.g. Martin et al. 2013, Malinsky et al. 2018, Meier et al. 2019, 2023). Although various modeling and simulation approaches have attempted to reveal the timing of past gene flow events, there remain serious challenges to accurately parameterize the models (Momigliano et al. 2021). Another option is to take advantage of the taxonomic diversity available in evolutionary radiations. By performing tests for introgression among the taxa in a radiation, it is possible to estimate the relative timing of introgression. Martin et al. (2013) used the diversity among *Heliconius* butterflies to demonstrate a history of continuous hybridization during the divergence of currently sympatric species pairs. Malinsky et al. (2018) expanded on this approach by performing tests for introgression between all sets of taxa in the Lake Malawi cichlid radiation, enabling researchers to evaluate which lineages showed evidence of hybridization and when it was likely to have occurred across the phylogeny of the group. They identified evidence of extensive ancient hybridization between the ancestors of modern cichlid lineages, as well as recent hybridization among closely related species found in similar ecological niches. These studies highlight how the taxonomic diversity present in evolutionary radiations can be used to identify the relative timing of past gene flow events. In this study, we expand on these previous examples to develop tests for the timing of introgression between island and mainland taxa in a radiation of monkeyflowers.

In addition to characterizing the evolutionary history of gene flow, understanding the fitness consequences of this hybridization can provide clues about the accumulation of reproductive isolation. In most cases, when functionally-relevant genetic variation is transferred into a foreign genetic background, it will decrease fitness, resulting in the removal of these alleles due to negative selection (Mallet 2005). These introgression-resistant loci are termed “barrier loci,” and as time proceeds, they should accumulate across the genome, leading to increased reproductive isolation (Wu 2001, Feder et al. 2012). However, positive selection following introgression can also result in the preservation and eventual fixation of these alleles (i.e. adaptive introgression) (Suarez-Gonzalez et al. 2018). For example, introgression of genetic material into a species that recently experienced a population bottleneck can introduce the necessary genetic variation to mitigate the negative effects of any deleterious genetic load that has accumulated (Schumer et al. 2018, Liu et al. 2022). Alternatively, the introgression of ecologically beneficial alleles can lead to adaptation, but this is expected to produce a more localized signature across the genome. By contrast, post-divergence gene flow can have no fitness consequences, in which case allele frequencies will be influenced primarily by genetic drift (Schumer et al. 2018).

To distinguish among these evolutionary processes, we expect certain genome-wide relationships to emerge. For example, the recombination rate determines how quickly introgressed alleles will be separated from resident alleles. Thus, we expect selection to rapidly remove deleterious, immigrant alleles in regions of low recombination where they remain linked with resident alleles. This will result in a positive relationship between introgression and recombination rate, provided the effects of reproductive isolation are widespread across the genome (Brandvain et al. 2014, Schumer et al. 2018, Martin et al. 2019). Similarly, because gene flow opposes divergence, we would expect a negative relationship between introgression and genetic divergence (Martin et al. 2013, Martin et al. 2019). In contrast, the repeated fixation of adaptively introgressed alleles will result in a genome-wide, negative relationship between introgression and recombination rate, such that longer, introgressed haplotypes are preserved in regions of low recombination (Duranton & Pool 2022, Feng et al. 2024). While this pattern is expected in scenarios where introgression acts to mitigate the widespread, deleterious effects of genetic load, introgression that results in local adaptation to a particular ecological environment will not lead to this genome-wide relationship. Finally, no relationship between introgression and recombination rate would imply the action of neutral processes (Schumer et al. 2018).

In this study, we take advantage of the diversity in the *Mimulus aurantiacus* species complex to investigate the evolutionary history and genomic consequences of introgression between a pair of taxa restricted to the Channel Islands off the coast of California. The *M. aurantiacus* complex is a radiation of seven closely related, woody shrub subspecies distributed throughout California that display extensive phenotypic variation in their floral and vegetative traits and diverged from their sister species roughly one million years ago (Tulig 2000, Tulig & Nesom 2012, Chase et al. 2017; Stankowski et al. 2019). However, despite extensive phenotypic differentiation, there is evidence of hybridization between many of the taxa (Streisfeld & Kohn 2005, Stankowski et al. 2019, Short & Streisfeld 2023). Indeed, ancient introgression resulted in the repeated evolution of red flowers, a trait that has been shown to contribute to both pollinator adaptation and speciation in this group (Stankowski & Streisfeld 2015, Short & Streisfeld 2023, Sobel & Streisfeld 2015, Stankowski et al. 2017). The Channel Islands are currently inhabited by two subspecies of *M. aurantiacus* that are known to hybridize (Wells 1980, Chase et al. 2017). The red-flowered subspecies *parviflorus* is endemic to many of the islands (Tulig 2000, Tulig & Nesom 2012) but is listed as rare by the California Native Plant Society (2023). Subspecies *longiflorus* has larger, yellow flowers and is found both on the Channel Islands and throughout mainland southern California (Tulig 2000, Tulig & Nesom 2012). Both taxa occur in close contact on the islands, where they tend to inhabit dry, hillside habitats in the chaparral (Beeks 1962).

We used whole genome sequence data from individuals sampled across the radiation to address three primary objectives. First, we developed an approach that incorporates multiple tests of introgression to estimate the history of hybridization among taxa and determine the relative timing of past gene flow events. Second, we assessed the genome-wide impacts of this introgression. Finally, we examined the relationship between introgression and recombination rate at different points throughout the history of the radiation to determine which processes have likely contributed to these patterns through time. Our results reveal that both recent and ancient introgression have shaped the genomic landscapes of these island taxa. Moreover, we found that the fitness effects of introgressed alleles can vary through time, such that the effects of selection against gene flow appear to increase with time, implying the accumulation of reproductive isolation.

## 2 MATERIALS AND METHODS

### 2.1 Genome sequencing and variant calling

Leaf tissue was collected from 27 samples from four locations on Santa Cruz Island, California, USA (Table S1), consisting of red-flowered *parviflorus*, yellow-flowered *longiflorus*, and their putative hybrids. Tissue was dried in silica in the field, and DNA was isolated using the Zymo Plant and Seed DNA kit following the manufacturer’s instructions. Sequencing libraries were prepared according to Gaio et al (2022), with slight modifications. Bead-Linked Transposase from the Illumina Nextera XT Kit was used for initial tagmentation, generating insert sizes in the range of 400 – 1200 bp. Multiplexed libraries were sequenced on the Illumina Novaseq 6000 using paired-end 150 bp reads at the University of Oregon’s Genomics Core Facility.

New sequences from Santa Cruz Island were combined with previously generated whole genome sequences from various *M. aurantiacus* subspecies from Stankowski et al. (2019) and Short & Streisfeld (2023), resulting in a final dataset containing 74 individuals. Raw reads were filtered using *fastp* to remove reads with uncalled bases or poor quality scores (Chen et al. 2018). The retained reads were then aligned to the reference assembly (Stankowski et al 2019) using *BWA* version 0.7.17 (Li & Durbin 2009). An average of 90.56% of reads aligned (range: 75.45% – 95.33%), and the average sequencing depth was 10× per individual (range: 6–18×). PCR duplicates were marked using *Picard* (https://broadinstitute.github.io/picard/). Variant calling was performed following Stankowski et al. (2019). We then phased the VCF using *BEAGLE* (Browning & Browning 2007), and further filtered the VCF file for biallelic SNPs using *vcftools* (Danecek et al. 2011). The final data set contained 12,749,566 SNPs across all 74 samples. Finally, we ran *UnifiedGenotyper* with the EMIT_ALL_CONFIDENT_SITES option to output all variant and invariant genotyped sites.

### 2.2 Admixture, PCA, and phylogenetic analysis

To determine how the island samples were related to the other taxa in the complex, we used *Admixture* (Alexander et al. 2009) to estimate ancestry proportions from all samples, with the number of clusters (K) set from 2 to 11. To further estimate ancestry among the admixed island samples and assess the relationship between samples collected on Santa Cruz Island and the mainland, we re-ran *Admixture* at K=2, but only using samples of *parviflorus*, *calycinus*, and Island and mainland *longiflorus*. Samples from the island that showed no evidence of admixture at K = 2 were used to assign individuals to subspecies. Admixed individuals were removed from further analysis, as they likely represented contemporary hybrids. To further assess clustering patterns among these samples, we also performed a principal components analysis (PCA) in *Plink* using the 31 samples from the island and the 7 mainland *longiflorus*.

To determine the phylogenetic relationships among samples from the island and mainland, we generated a maximum likelihood consensus tree using *IQ-TREE* v1.6.12 (Nguyen et al. 2015). We used a concatenated dataset consisting of 12,749,566 biallelic SNPs that did not include island samples that showed evidence of admixture in the above analyses (resulting in 63 individuals, see Results).

### 2.3 Tests for the timing of admixture

To test for genome wide evidence of introgression, we used *dsuite* (Malinsky et al. 2021) to calculate Patterson’s D (Green et al. 2010) and the *f4*-ratio (Reich et al. 2009) for all possible trios of ingroup taxa, using *M. clevelandii* as the outgroup. Patterson’s D and the *f4*-ratio measure asymmetries in the numbers of sites with ABBA and BABA patterns (where A and B are ancestral and derived alleles, respectively) across a phylogeny with three ingroup taxa and an outgroup that have the relationship (((P1, P2), P3) O). A significant excess of either pattern gives a nonzero value of D, which is taken as evidence that gene flow has occurred between P3 and one of the sister taxa. We then calculated the fbranch statistic to identify recent and historical signatures of hybridization (Malinsky et al. 2018, 2021). Given a set of *f4*-ratios calculated for all possible trios among a set of closely related taxa, *fbranch* uses the inferred phylogenetic relationship among these taxa, as well as variation in the phylogenetic distance between the sister taxa used as P1 and P2, to assign introgression to specific branches on a phylogenetic tree.

To estimate how introgression varied across the genome, we calculated the admixture proportion (*f_d_*) (Martin et al 2015) in 100 kb non-overlapping windows using the *ABBABABAwindows.py* Python script (https://github.com/simonhmartin/genomics_general). Similar to Patterson’s D and the *f4*-ratio, *f_d_* searches for asymmetries in the number of ABBA and BABA sites across the genome, but it has been optimized specifically for use in genomic windows.

In addition to examining how *f_d_* varies across the genome, we took advantage of the diversity of taxa in the *M. aurantiacus* species complex to estimate the relative timing of introgression between the taxa on Santa Cruz Island (Fig 1). Specifically, alleles that were introgressed into the common ancestor of Island and mainland *longiflorus* should be present in roughly equal proportions in both descendent lineages, making it difficult for *f_d_* to distinguish shared ancestral variation from introgression. However, by gradually increasing the phylogenetic distance between the two sister taxa used in these tests, we should be able to identify introgression that occurred at various points further back in time (Martin et al 2013; Short and Streisfeld 2023). This can be identified as an increase in mean *f_d_* with levels of sequence divergence. Alternatively, if introgression occurred only after the divergence between Island and mainland *longiflorus*, then we would not expect *f_d_* to increase with levels of sequence divergence.

**Figure 1.**
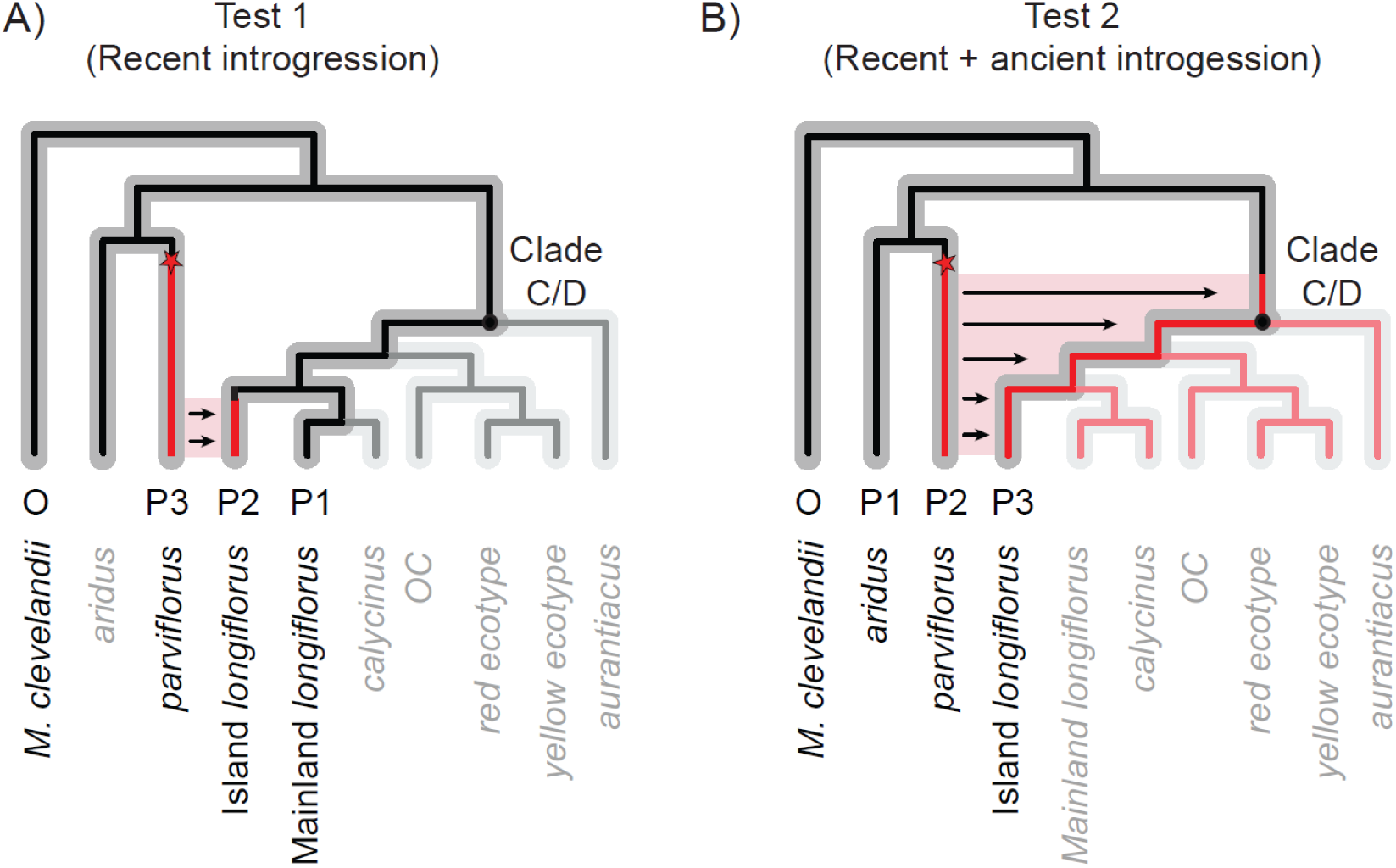
Evolutionary radiations can be used to estimate the relative timing of introgression. (A-B) In a four-taxon tree, with three ingroup taxa (P1–P3) and an outgroup (O), introgression between P2 and P3 only can be identified if gene flow occurred after the split between the two sister taxa (P1 and P2). The species tree from the *M. aurantiacus* radiation is presented in gray, with tips grayed out to show only the four taxa used in the tests for introgression (denoted by P1–P3, and O). The ancestral (black) and derived (red) alleles at a locus are indicated as lines. Mutations are indicated by red stars. By using different pairs of taxa with increasing levels of divergence from one another as P1 and P2, we can track introgression that occurred further back in time. (A) To estimate recent introgression (i.e., the genetic variation that was introgressed after the divergence of Island and mainland *longiflorus*), we performed Test 1, which includes mainland *longiflorus* as P1, Island *longiflorus* as P2, and *parviflorus* as P3. (B) To estimate recent ancient introgression (i.e., any genetic variation that was exchanged between the ancestor of *parviflorus* and the common ancestor of clades C and D), we performed Test 2, which includes the more distantly related *aridus* and *parviflorus* as P1 and P2, and Island *longiflorus* as P3. The tests performed do not specify the direction of introgression. For simplicity, the examples shown in the figure do indicate directionality, but the same conclusion would be drawn about the timing of introgression if the direction was reversed

To examine this, we ran two distinct tests for introgression by varying the set of taxa used in each test (Fig 1). To identify the potential for recent introgression, we calculated mean *f_d_* among 100 kb windows, setting Island *longiflorus* as P2, *parviflorus* as P3, and varying the taxon used as P1. Hereafter, we refer to these as Test 1. Then, to determine whether there was evidence for recent and ancient introgression, we took advantage of the more extensive divergence between *parviflorus* and *aridus*, which occurred prior to the split between taxa in clades C and D (Stankowski et al 2019). If introgression predated the divergence between Island and mainland *longiflorus*, then we would expect *f_d_* to increase when these more distantly related taxa were included as P1 and P2. Thus, we set *aridus* as P1, *parviflorus* as P2, and Island *longiflorus* as P3. These analyses will be referred to as Test 2. *M. clevelandii* was used as the outgroup for all calculations of *f_d_*.

To further demonstrate whether there was recent introgression between *parviflorus* and Island *longiflorus,* we re-calculated Test 2 to obtain the mean *f_d_* among windows using *aridus* as P1, *parviflorus* as P2, and the various taxa from clades C and D as P3. Among these tests, if mean *f_d_* is greatest when Island *longiflorus* is P3, this would indicate both ancient introgression that predated the divergence of clades C and D, as well as recent introgression between Island *longiflorus* and *parviflorus*. However, if introgression occurred recently between *parviflorus* and any of the other taxa in clades C and D, then we would expect *f_d_* to be equal to or greater than the *f_d_* value calculated using Island *longiflorus* as P3.

To test for significant differences among the mean *f_d_* values, we fit linear models, with *f_d_* as the dependent variable and sequence divergence (da) between P1 and P2 as the independent variable. We then used the *emmeans* package in R (Lenth, 2019) to perform pairwise comparisons of the estimated marginal mean *f_d_* values from the linear model. To estimate levels of sequence divergence, we calculated da, which describes the difference in divergence between taxa (*d_xy_*) relative to the mean diversity within taxa (π). We calculated *d_xy_* and π in 100 kb windows using *PIXY* version 1.2.5 (Korunes & Samuk, 2021), with both variant and invariant sites included.

### 2.4 The relationship between genome wide phylogenetic discordance and introgression

To further describe the history of introgression with *parviflorus*, we explored patterns of phylogenetic discordance across the genome using *TWISST* (Martin & Van Belleghem 2017). Given a set of samples from multiple ingroup taxa, *TWISST* builds trees in genomic windows and then calculates the support for that topology among all possible topologies, which is referred to as the topology weighting. If all samples from each taxon are reciprocally monophyletic, then that topology is given a weighting of 1.0 for that window. Topological discordance among samples within a taxon will result in lower topology weightings for that window. We used *TWISST* to identify variation in the relationships among *aridus*, *parviflorus*, Island *longiflorus*, and mainland *longiflorus* in 100 kb genomic windows, with *M. clevelandii* as the outgroup. With four ingroup taxa, this resulted in 15 possible topologies, which were then partitioned into those that support: *a*) the species tree, where *aridus* and *parviflorus* were sister and reciprocally monophyletic relative to Island *longiflorus* and mainland *longiflorus*; *b*) the introgression tree, where *parviflorus* and Island *longiflorus* were sister to one another relative to *aridus* and mainland *longiflorus*; and *c*) the ancient introgression tree, where *parviflorus* was sister to mainland and Island *longiflorus* relative to *aridus*. By quantifying variation in the support for these topologies across the genome, we can identify how often ancient and recent introgression contributed to phylogenetic discordance.

However, topological discordance can also be influenced by incomplete lineage sorting. Thus, to confirm that regions with high support for the introgression or ancient introgression trees were consistent with admixture, we first identified 100 kb windows with topology weightings of 1.0 for the species tree, introgression tree, or the ancient introgression tree, and then investigated the distribution of *f_d_* values within these windows. We calculated *f_d_* from Test 1 (with mainland *longiflorus* as P1) to identify the effects of recent introgression, while *f_d_* from Test 2 (with Island *longiflorus* as P3) was used to identify the combined effects of recent and ancient introgression. By taking the difference between these two sets of *f_d_* values, we should be able to identify only the signal of ancient introgression, which we refer to as “ancient *f_d_*.” For example, when mainland and Island *longiflorus* are set as P1 and P2 in Test 1, we can only identify introgression with *parviflorus* that has occurred since they diverged from each other (Fig 1). However, when *aridus* and *parviflorus* are set as P1 and P2 in Test 2, the more ancient divergence between them allows us to identify older introgression with the ancestor of *parviflorus* that occurred prior to the split between clades C and D, as well as recent introgression that occurred after the divergence between mainland and Island *longiflorus*. Thus, the difference between the two provides us with the signal of ancient introgression that occurred prior to the split of clades C and D. We determined how ancient and recent introgression impacted the relationships among these taxa by quantifying the topology weightings for the species tree, introgression tree, and ancient introgression tree within the top 5% of *f_d_* windows.

### 2.5 The genomic consequences of introgression

Genome wide heterogeneity in admixture can be caused by one (or a combination) of several processes: selection against gene flow, adaptive introgression, or genetic drift. To determine which of these processes may be contributing to genome wide variation in introgression, we estimated the relationship between the recombination rate, relative genetic divergence (*F_ST_*), and *f_d_* due to both recent and ancient introgression. Under a model of polygenic selection against gene flow, we expect a negative relationship between *f_d_* and *F_ST_*, but a positive relationship between *f_d_* and recombination rate (Brandvain et al. 2014, Schumer et al. 2018, Liu et al. 2022). Alternatively, if there is a genome-wide signal of adaptive introgression, which could occur to mitigate the effects of deleterious genetic load induced by hybridization (Feng et al. 2024), then we would expect a negative relationship between *f_d_* and recombination rate (Duranton & Pool 2022). In the absence of selection, we expect no relationship between *f_d_* and recombination rate. Finally, as reproductive isolation accumulates through time, we expect a stronger positive relationship between recombination and *f_d_* for estimates of recent introgression but a weaker relationship with ancient introgression. This is because fewer reproductive barriers would be in place if gene flow occurred deeper in the past.

*F_ST_* was estimated between *parviflorus* and Island *longiflorus* in 100 kb non-overlapping windows using the *popgenwindows.py* Python script (https://github.com/simonhmartin/genomics_general), with only variant sites included. *f_d_* values from Test 1, Test 2, and “ancient *f_d_*” were correlated with *F_ST_* using Spearman’s correlation coefficient in R. To estimate the relationship between *f_d_* and recombination rate, we partitioned the *f_d_* values calculated in 100kb windows into quantile bins of recombination rates that were calculated previously in 500 kb windows by Stankowski et al. (2019). We then calculated the mean and 95% confidence intervals for *f_d_* within each recombination rate quantile bin and fit a linear model with *f_d_* as the dependent variable and the recombination rate quantile bins as the independent variable. We then used the *emmeans* package (Lenth, 2019) to perform pairwise comparisons of the estimated marginal mean *f_d_* values for the different quantile bins from the linear model.

It has also been hypothesized that introgression between closely related island endemic taxa may contribute to the maintenance of high levels of genetic diversity among island taxa that are geographically isolated from other populations of the same taxon (Carlquist 1966). If differences in genetic load between mainland and island endemics affect the outcomes of introgression between these taxa, we would expect to find reduced genetic diversity and a lower effective population size in the island taxa as compared to their mainland relatives (Schumer et al. 2018, Liu et al. 2022). To determine whether there is evidence of reduced genetic diversity among the hybridizing island taxa, we asked whether the mean π in both island subspecies differed among the other taxa in the complex. To identify differences in the effective population sizes of these species, we used PSMC (Li and Durbin 2011) to estimate the effective population size through time for all samples from the island taxa and their most closely related mainland relatives. In accordance with Stankowski et al. (2023), we assumed a generation time of 2 years and a mutation rate of 7×10^-9^ for these calculations.

### 2.6 Identifying signatures of adaptive introgression

Extensive heterogeneity in admixture across the genome raises the possibility that adaptive introgression has maintained foreign ancestry in particular regions. To determine whether regions of elevated *f_d_* display signatures of adaptive introgression between *parviflorus* and island *longiflorus*, we calculated the frequency of Q95 sites found in windows across the genome (Racimo et al. 2016, Feng et al. 2024). Q95 sites are defined as those that are fixed (at a frequency of 1.0) in the donor taxon, near fixation (at a frequency greater than the 95^th^ percentile) in the recipient taxon, and absent in the taxon that is sister to the recipient. Thus, any derived alleles that are present at high frequency in a recipient taxon that are also absent in the unadmixed sister taxon but are fixed in a more distantly related donor taxon, are candidates for adaptive introgression. Genomic regions that display an excess of these putatively adaptively introgressed alleles are even more likely to have been targets of positive selection, because selection will have correlated effects on linked sites. Thus, by identifying an elevated frequency of Q95 sites in 100 kb windows across the genome, we should be able to identify genomic regions that have been adaptively introgressed.

To identify Q95 sites, we first identified all derived SNPs that were polymorphic among samples. Derived SNPs were defined as those that were fixed in *M. clevelandii* and variant in at least one chromosome from samples of mainland *longiflorus*, Island *longiflorus*, *parviflorus*, or *aridus*. We then determined the number of derived sites that were fixed in *parviflorus*, absent in mainland *longiflorus*, and present in at least one haplotype in Island *longiflorus*. These represented sites that were introgressed from *parviflorus* into Island *longiflorus*. The 95th percentile of the allele frequency distribution among introgressed sites in Island *longiflorus* (=0.875) was used as a cutoff to identify adaptively introgressed sites (i.e., Q95 sites) (Racimo et al. 2016, Feng et al. 2024). For each 100 kb window, we then divided the number of Q95 sites by the total number of derived SNPs that were present on at least one chromosome in Island *longiflorus* to estimate the frequency of Q95 sites in each window.

To identify signatures of adaptive introgression that were transferred from Island *longiflorus* into *parviflorus*, we identified derived SNPs that were fixed in Island *longiflorus*, present on at least one haplotype in *parviflorus*, but were absent in mainland *longiflorus*. In this case, the 95^th^ percentile of the allele frequency distribution in *parviflorus* included only sites fixed in *parviflorus*. We then divided the number of Q95 sites by the total number of derived SNPs that were present on at least one chromosome in *parviflorus* in 100 kb windows.

To further explore signatures of adaptive introgression for the region with the highest frequency of Q95 sites across the genome, we calculated *f_d_*, *d_xy_*, *F_ST_*, π, and the frequency of Q95 sites in 10 kb windows with 1000 bp steps (see Results).

## 3 RESULTS

### 3.1 Evolutionary relationships and hybridization on Santa Cruz Island

At K=8, *Admixture* assigned samples to ancestry groups that were consistent with previous analyses of the phylogenetic history of these subspecies (Fig 2A, Fig S1) (Chase et al. 2017, Stankowski et al. 2019, Short and Streisfeld 2023). The newly sequenced Santa Cruz Island samples contained a mixture of individuals with both pure and admixed ancestry, suggesting the occurrence of ongoing hybridization between *parviflorus* and Island *longiflorus* (Fig 2A). By re-running Admixture at K=2, including only the island samples, as well as mainland *longiflorus* and *calcyinus,* we identified 12 “unadmixed” *parviflorus* and 8 “unadmixed” Island *longiflorus* (with q-scores greater than 0.99; Fig 2B). We also identified 11 admixed samples from Santa Cruz Island, which were removed from subsequent analyses. Results from the PCA were largely consistent with Admixture, with “unadmixed” Island *longiflorus* and *parviflorus* samples clustering separately along PC1 and admixed samples distributed between them (Fig 2C).

**Figure 2.**
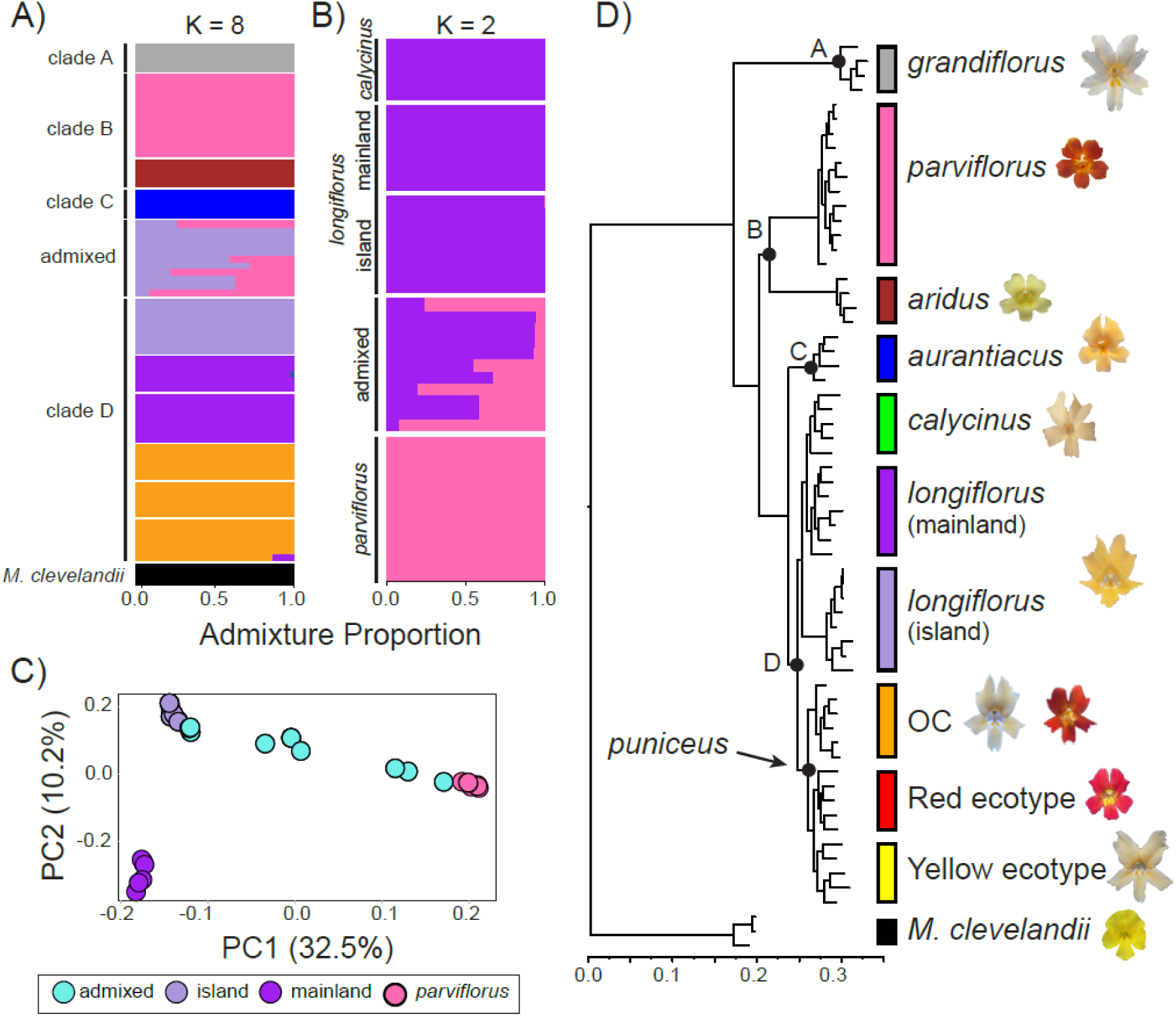
The relationships among the taxa in the *M. aurantiacus* species complex. (A) The ancestry proportions from *Admixture* at K = 8 for all the subspecies and their sister species, *M. clevelandii*. (B) Ancestry proportions at K = 2 from a run that included only the island samples, mainland *longiflorus*, and *calycinus*. (C) Plot of the first two principal components from the island samples and mainland *longiflorus*. The percent variation explained by each axis is reported. (D) Phylogenetic tree showing evolutionary relationships, with representative photographs of each taxon’s flower. The four major clades of the radiation are labelled with black letters. The tips corresponding to each taxon are indicated by a colored bar.

The consensus species tree was consistent with previous analyses that revealed four primary clades (Stankowski and Streisfeld 2015, Chase et al. 2017, Short & Streisfeld 2023). Clade A consisted entirely of subspecies *grandiflorus*, clade B showed *parviflorus* and *aridus* as sister taxa, clade C consisted entirely of *aurantiacus*, and the remaining taxa from southern California comprised the diverse clade D (Fig 2D). Within clade D, Island *longiflorus* was sister to both mainland *longiflorus* and *calycinus*. To confirm this relationship, an additional run of Admixture was performed at K=3 (Fig S2), revealing three distinct ancestry groups that corresponded to Island *longiflorus*, mainland *longiflorus*, and *calycinus*.

### 3.2 Evidence of recent and ancient hybridization

Consistent with Stankowski et al (2019), our calculations of the *fbranch* statistic revealed evidence of widespread introgression, particularly among the very recently diverged taxa within clades C and D (Fig 3A). In addition, we identified evidence of hybridization between *aurantiacus* and *grandiflorus,* as well as between *aridus* and the clade D taxa, confirming previous results by Short and Streisfeld (2023). We observed a strong signal of introgression between *parviflorus* and all the taxa from clades C and D, but this signal was the strongest between *parviflorus* and Island *longiflorus* (Fig 3A). This raises the possibility that there may have been recent introgression between *parviflorus* and Island *longiflorus*, as well as historical hybridization between *parviflorus* and the ancestor of clades C and D.

**Figure 3.**
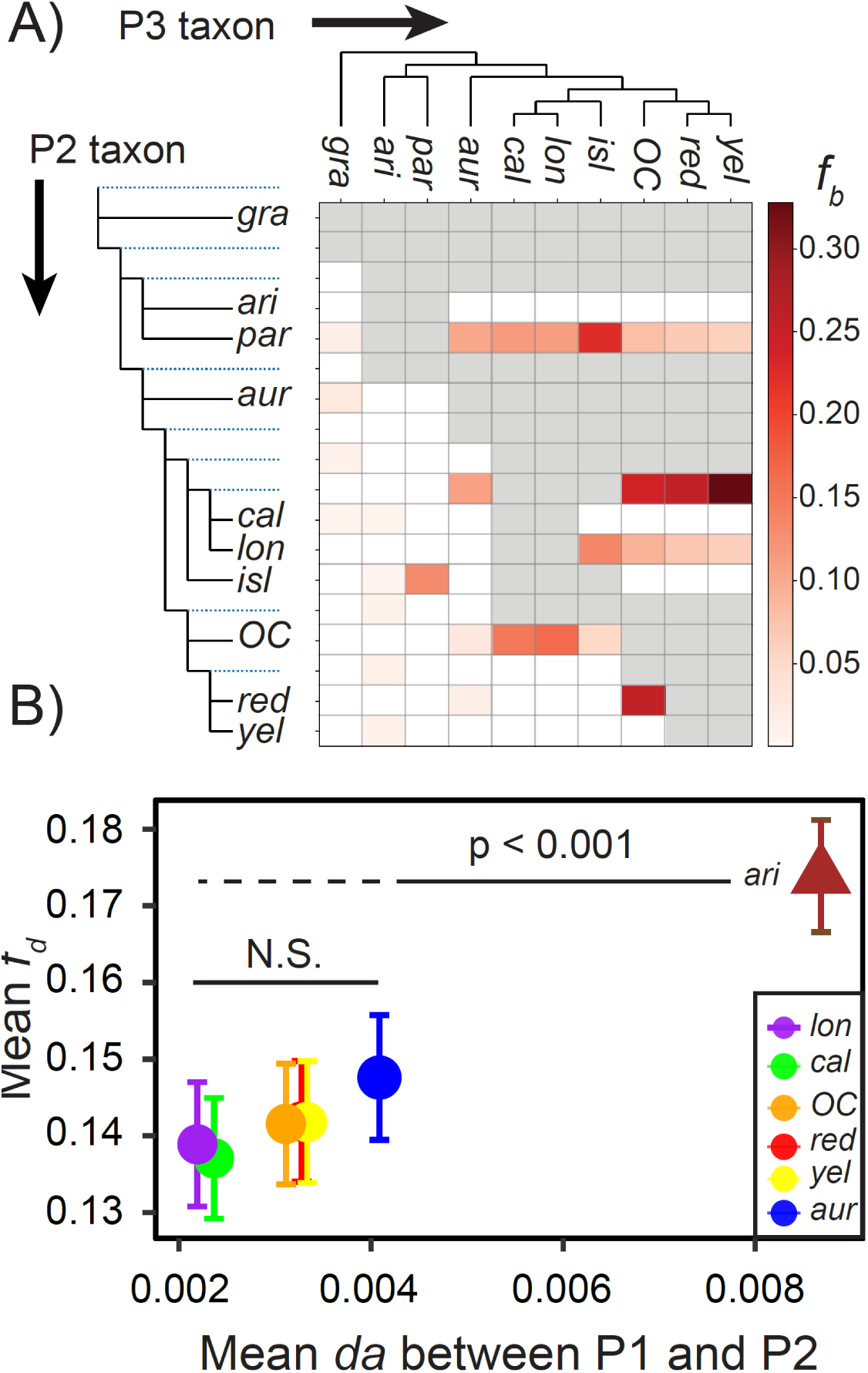
Hybridization history in the *Mimulus aurantiacus* species complex. (A) Genome-wide patterns of introgression using the f-branch statistic (fb). This analysis uses knowledge of the relationships among taxa and the results of the *f4*-ratio (calculated using all possible trios of ingroup taxa) to estimate the relative position of hybridization on a species tree. The color gradient represents the values of fb, with darker colors indicating greater evidence of introgression. (B) Mean and 95% confidence intervals for Test 1and Test 2 *fd* values calculated in 100 kb windows are plotted against mean sequence divergence (da) between the taxa used as P1 and P2 for the calculation of *fd*. Test 1 calculations of *fd* were performed using the various clade C and D taxa as P1, Island *longiflorus* as P2, and *parviflorus* as P3. Test 2 calculations of *fd* were performed using *aridus* as P1, *parviflorus* as P2, and Island *longiflorus* as P3. Colors indicate the P1 taxon used to calculate *fd* for the specific test, with mainland *longiflorus* in purple (*lon*), *calycinus* in green (*cal*), Orange County *puniceus* in orange (*OC*), the red ecotype in red (*red*), the yellow ecotype in yellow (*yel*), *aurantiacus* in blue (*aur*), and *aridus* in brown (*ari*). Shapes indicate which test was performed, with circles indicating Test 1 and triangles indicating Test 2.

To test these hypotheses, we performed calculations of *f_d_* using taxa at various levels of sequence divergence from one another. This allowed us to identify recent and historical signals of introgression (Fig 1). From Test 1, we identified no significant increase in mean *f_d_* with sequence divergence (da) when the different taxa from clades C and D were set as P1 (Fig 3B, Table S2). This indicates introgression was recent and occurred only after the split between Island and mainland *longiflorus*. However, when the more diverged *aridus* and *parviflorus* were set as P1 and P2 in Test 2, we found a significantly higher mean *f_d_* (Fig 3B, Table S2), indicating both recent hybridization between Island *longiflorus* and *parviflorus*, as well as ancient hybridization between the ancestor of modern *parviflorus* and the ancestor of clades C and D. In addition, mean *f_d_* was greatest when island *longiflorus* was set as P3 (Fig S3, Table S3), confirming that Test 2 identified the signals of both recent and ancient introgression. In addition to the presence of ongoing hybridization between the island taxa, these results reveal that introgression with *parviflorus* has occurred at two time points, once between its ancestor and the ancestor of clades C and D, and then again, in the more recent past with Island *longiflorus*.

### 3.3 Ancient and recent hybridization contribute to genome wide phylogenetic discordance

Using *TWISST*, we defined tree topologies that supported the ‘species tree,’ a set of ‘introgression trees’ where Island *longiflorus* and *parviflorus* were sister, and an ‘ancient introgression’ topology where *parviflorus* was sister to both mainland and Island *longiflorus* (Fig 4A). By scanning the genome for the distribution of these tree topologies, we found widespread evidence of phylogenetic discordance (Fig 4B). We identified 553 windows with topology weightings of 1.0 for the species tree, 61 windows that supported the introgression tree, and 419 trees that matched the ancient introgression tree (Fig 4C) (a remaining 915 windows did not match any of these topologies or had topology weightings less than 1.0, which we refer to as ‘unresolved’). Despite there being nearly as many topologies that supported a history of introgression as the species tree, this phylogenetic discordance also may have been caused by incomplete lineage sorting.

**Figure 4.**
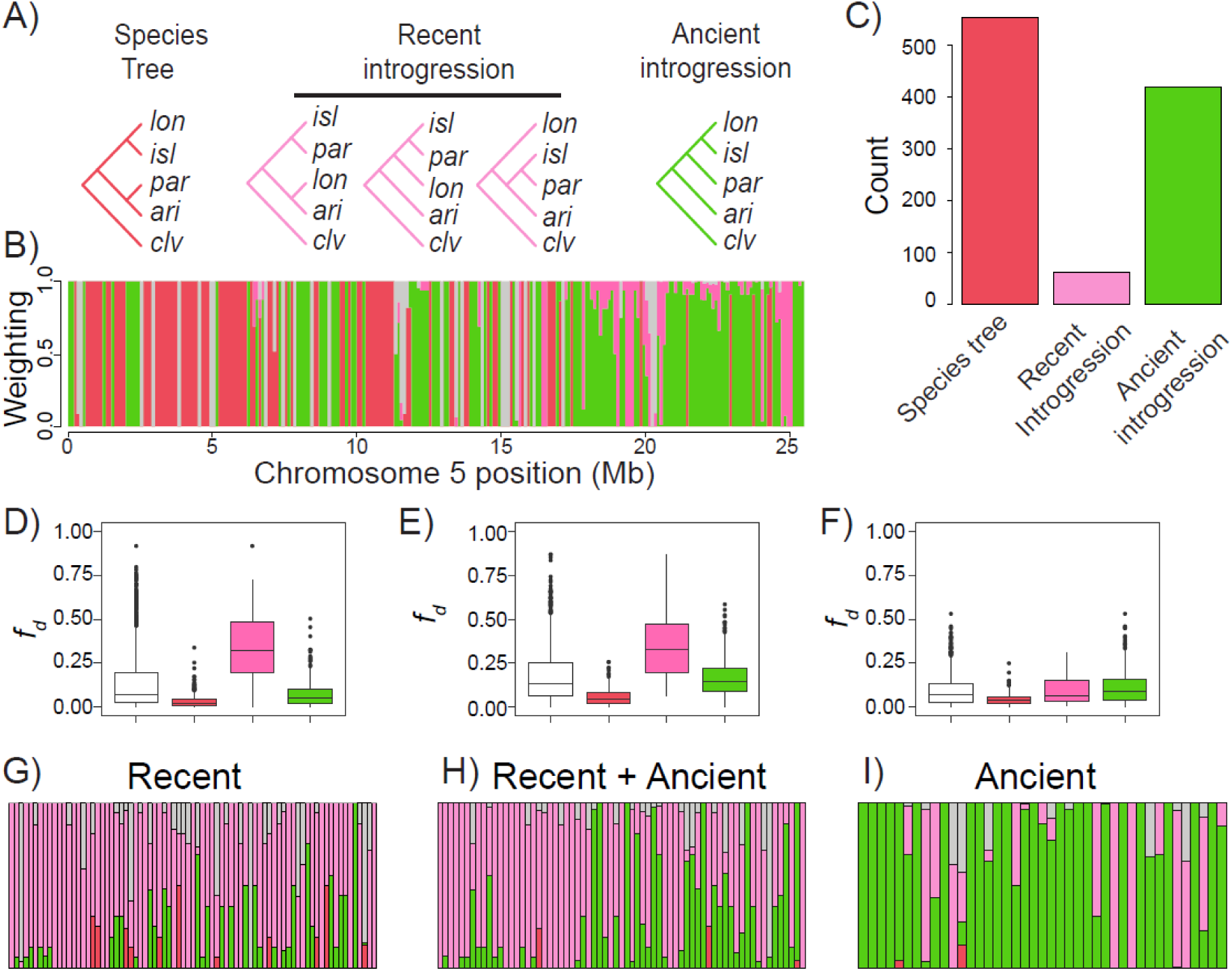
Genome wide variation in levels of phylogenetic discordance. (A) Topologies that represent the species tree (red), recent introgression trees (pink), and the ancient introgression tree (green). (B) Representative variation in topology weighting across chromosome 5. Red bars indicate support for the species tree, pink bars indicate support for the recent introgression tree, green bars indicate support for the ancient introgression tree, and gray bars indicate no support for these topologies. (C) The number of 100 kb windows across the genome containing a topology weighting of 1.0 for the species tree, recent introgression tree, and the ancient introgression tree. (D-F) Distribution of *fd* values among windows with a topology weighting of 1.0 for either the species tree (red), recent introgression tree (pink), or the ancient introgression tree (green). The distribution for the entire genome is provided in white. (D) Test 1 *fd* values. (E) Test 2 *fd* values. (F) Ancient *fd* values, calculated by taking the difference between Test 2 and Test 1 *fd* values. (G-I) Distribution of topology weightings among the 5% of windows with the highest *fd* values. (G) Test 1 *fd* values. (H) Test 2 *fd* values. (I) Ancient *fd* values.

To determine if discordance could be attributed to introgression or incomplete lineage sorting, we tested for an association between these topologies and *f_d_* across the genome. Consistent with a primary role for introgression, we found that windows supporting the ‘introgression tree’ displayed higher Test 1 and Test 2 *f_d_* values than the other topologies (Figs 4D and 4E). However, there were some windows with high support for the ‘introgression tree’ that had low values of *f_d_*, suggesting that some of the discordance may have been caused by incomplete lineage sorting. In windows with complete support for the ‘ancient introgression tree,’ *f_d_* values from Test 2 were higher than for Test 1 (compare Fig 4D and 4E), again consistent with Test 2 identifying signals of both recent and ancient introgression. When we calculated the difference between *f_d_* values from Test 2 and Test 1, the windows with the highest ‘ancient *f_d_*’ values largely corresponded to those with complete support for the ‘ancient introgression’ topology (Fig 4F). By focusing only on the top 5% of windows with the highest *f_d_* for both Test 1 and Test 2, we found greater support for the ‘introgression’ and ‘ancient introgression’ topologies (Figs 4G-H), with more evidence of ancient introgression for Test 2. This pattern became even more clear when we plotted the distribution of topologies for the top 5% of ‘ancient *f_d_*’ vales (Fig 4I). Thus, while there is some evidence for incomplete lineage sorting, much of the phylogenetic discordance appears to be caused by recent and ancient introgression.

### 3.4 The genomic consequences of recent and ancient hybridization

We identified extensive variation in the extent of introgression across the genome (Fig 5, S4). Raw values of *f_d_* in 100 kb windows ranged from 0.0 to around 0.9 (Fig S4). To determine the evolutionary processes responsible for this extreme heterogeneity in introgression across the genome, we compared *f_d_* with levels of genetic divergence and recombination rates. We identified a strong negative relationship between *f_d_* and *F_ST_* for both Test 1 and Test 2 (r > −0.70; Fig. 5D, 5E), implying that negative selection has likely purged deleterious foreign genetic variation across the genome. By contrast, although the relationship between *F_ST_* and *f_d_* remained negative when we examined the impacts of ancient introgression, it was considerably weaker (r = −0.26, Fig 5F).

**Figure 5.**
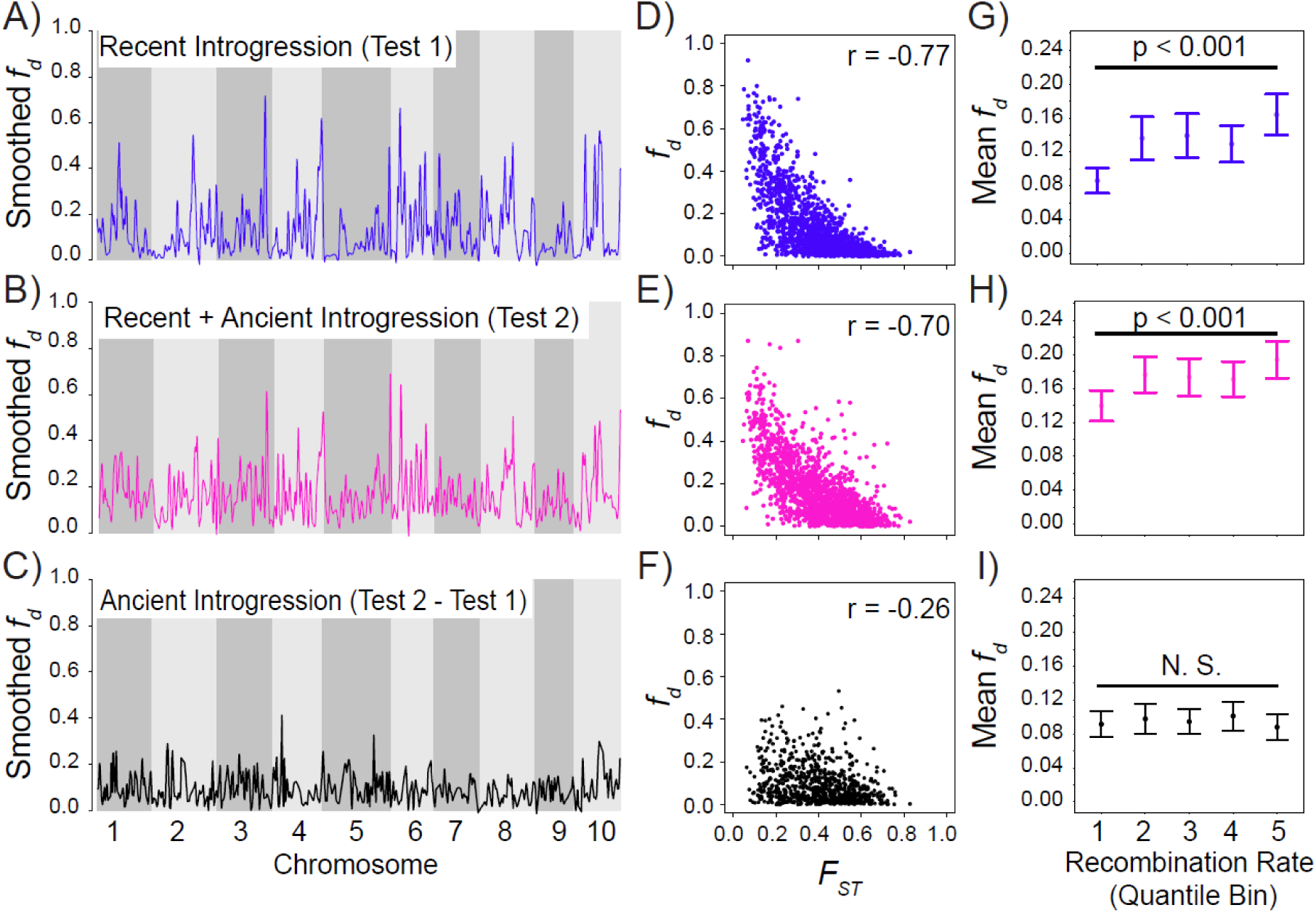
Genome wide variation in introgression. (A-C) Loess-smoothed *f_d_* values plotted across the 10 chromosomes of the *M. aurantiacus* genome. Colors indicate how the *f_d_* values were calculated: (A) Test 1 *f_d_* values, (B) Test 2 *f_d_* values, (C) Ancient *f_d_*. (D-F) Scatterplots showing the relationship between *f_d_* and genetic differentiation (*F_ST_*) between Island *longiflorus* and *parviflorus*, with the correlation coefficient between the statistics in the upper right hand corner of each plot. (D) The relationship between recent introgression (i.e., Test 1 *f_d_*) and *F_ST_*. (E) The relationship between both recent and ancient introgression (i.e., Test 2 *f_d_*) and *F_ST_*. (F) The relationship between ancient introgression and *F_ST_*. (G-I) Mean and 95% confidence intervals of *f_d_* calculated among quantile bins of recombination rate. Quantile bins of recombination rate are as follows: 1 = 0–0.824 cM/Mb; 2 = 0.824–1.51 cM/Mb; 3 = 1.51–2.33 cM/Mb; 4 = 2.33–3.66 cM/Mb; 5 = 3.66–10.9 cM/Mb. (G) Test 1 *f_d_* values, (H) Test 2 *f_d_* values, and (I) ancient *f_d_* values.

We found an overall positive relationship between recombination rate and mean *f_d_* for Tests 1 and 2 (Fig 5G, 5H). Specifically, the recombination rate quantile bin with the lowest recombination (0–0.824 cM/Mb) had significantly lower mean *f_d_* values than the highest recombination quantile (3.66–10.9 cM/Mb) for both Tests 1 and 2 (Table S4, S5). In addition, for Test 1, mean *f_d_* in the lowest recombination rate quantile was significantly lower than all four of the remaining higher quantiles, but this was not the case for Test 2, which showed no significant differences in mean *f_d_* among intermediate recombination quantile bins. The slightly weaker relationship for Test 2 suggests the possibility that the effects of selection may have varied through time. Consistent with this hypothesis, we identified no significant difference in ancient *f_d_* values between any of the recombination rate bins (Figure 5I, Table S6).

We observed no difference in average nucleotide diversity (π) among taxa, including the island endemics (Fig S5). Furthermore, we found no evidence for recent or historical reductions in effective population size in the island taxa (Fig S6). However, we did observe a small increase in effective population size in *parviflorus* and Island *longiflorus* around 100,000 years ago, suggesting that hybridization between these subspecies may have started around that time.

### 3.5 Evidence of localized adaptive introgression

To investigate evidence for adaptive introgression in regions of elevated *f_d_*, we calculated the frequency of Q95 sites in 100 kb windows across the genome (Fig S7). As might be expected, most windows displayed no increases in the frequency of Q95 sites. However, we did identify clear pileups of elevated Q95 ratios in a few regions (Fig S7), with one region in particular showing more than 40% of derived sites within two adjacent windows that matched the Q95 pattern (chromosome 5, 24.7-24.9 Mbp). These same windows had *f_d_* values greater than 0.85 for Test 2 and about 0.7 for Test 1 (Fig S4). Such an elevated local signal of introgression surrounded by regions of considerably higher admixture suggests the possibility of adaptive introgression. In addition, although we found an overall positive relationship between *f_d_* and π, these two windows have the highest *f_d_* for both Tests 1 and 2 but some of the lowest nucleotide diversity across the genome (Fig S8).

By zooming in on this region on chromosome 5, we identified a strong signal of introgression that spanned approximately 300 kb (Fig 6A). Furthermore, we also identified reduced *d_xy_* and *F_ST_* between *parviflorus* and Island *longiflorus* and a clear increase in the frequency of Q95 sites transferred from Island *longiflorus* into *parviflorus* (Fig 6). In contrast, we did not see a signal of elevated Q95 when considering introgression in the other direction. In addition, we also identified elevated *F_ST_* and *d_xy_* between *parviflorus* and *aridus*, but no clear increase in *F_ST_* or *d_xy_* between Island and mainland *longiflorus* (Figure 6B, 6C), providing further support that adaptive introgression in this region occurred from Island *longiflorus* into *parviflorus*. However, we observed no clear decrease in π for either Island taxon (Figure 6D).

**Figure 6.**
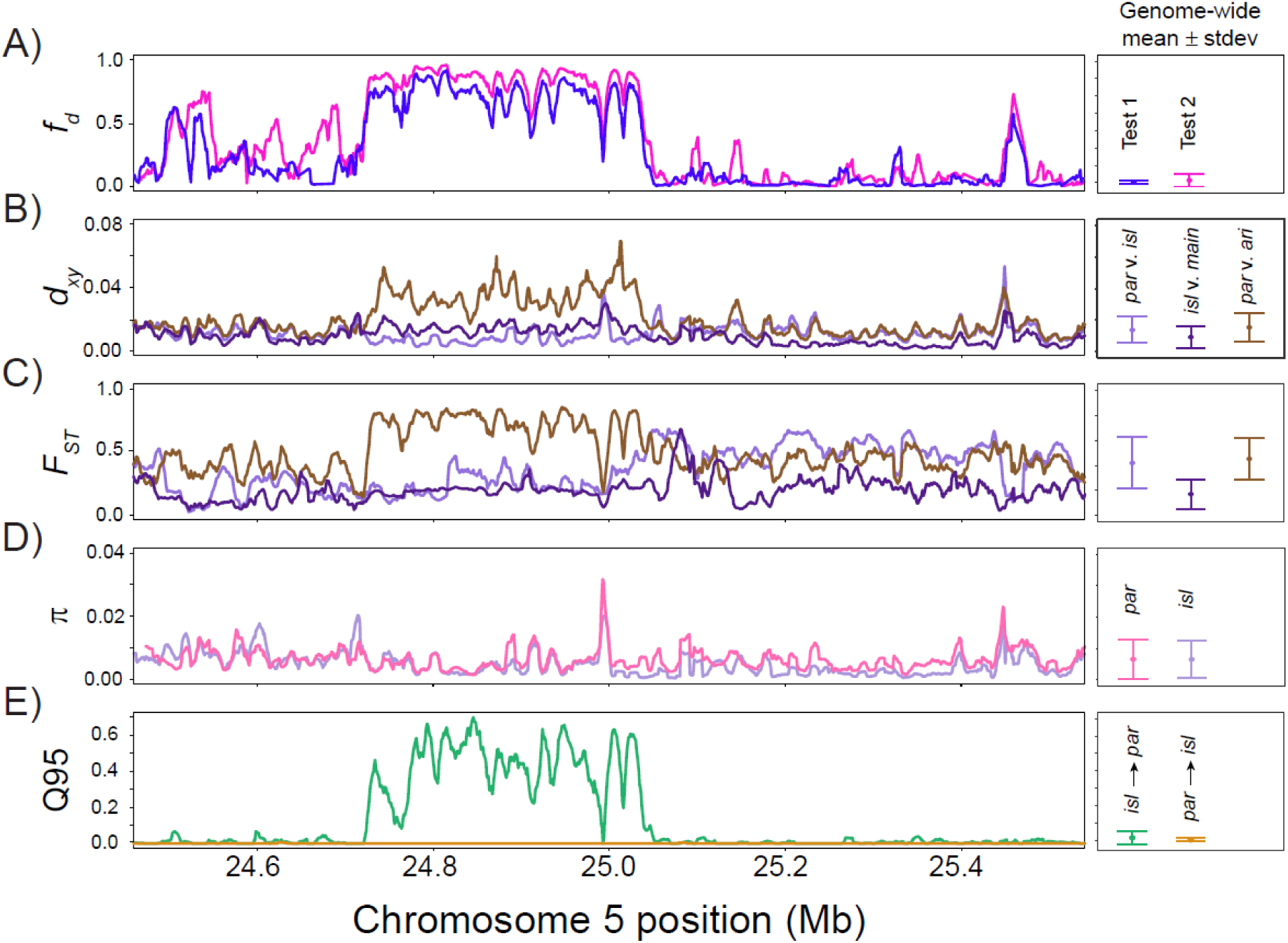
Hybridization leads to adaptive introgression from Island *longiflorus* into *parviflorus*. On the left, the admixture proportion (*f_d_*), genetic divergence (*d_xy_*), genetic differentiation (*F_ST_*), genetic diversity (π), and Q95 are plotted across an 800 kb region on chromosome 5 identified as a candidate region using the Q95 statistic (Fig S7). On the right of each plot is the genome-wide mean ± standard deviation of each statistic, calculated in 10 kb windows and 1000 bp steps across all 10 chromosomes. (A) Both Test 1 and Test 2 *f_d_* values indicate a defined region of elevated introgression. (B) *d_xy_* between *parviflorus* and *aridus* (brown) is clearly elevated across this region. However, *d_xy_* between *parviflorus* and Island *longiflorus* (light purple) and *d_xy_* between Island and mainland *longiflorus* (purple) are nearly equal to genome wide levels. (C) *F_ST_* between *parviflorus* and Island *longfilorus* (light purple) is reduced in this region, and *F_ST_* between Island and mainland *longiflorus* (purple) is nearly equal to genome wide levels. *F_ST_* between *parviflorus* and *aridus* (brown) displays a clear increase in this region as compared to genome wide levels. (D) No clear decrease in π was observed for either *parviflorus* (pink) or Island *longiflorus* (light purple). (E) A clear increase in the number of alleles that are shared between Island *longiflorus* and *parviflorus* but absent in *aridus* (green) was observed within this region. In contrast, no increase in the number of alleles that are shared between *parviflorus* and Island *longiflorus* but absent in mainland *longilforus* (yellow) was observed within this region, indicating unidirectional introgression from Island *longiflorus* into *parviflorus*.

## 4 Discussion

Hybridization can occur throughout the history of a radiation, but the evolutionary consequences of this mixing can vary over time. In this study, we took advantage of the diversity of taxa present in the *Mimulus aurantiacus* species complex to demonstrate that introgressive hybridization occurred at different points during the history of the group. We identified evidence of recent introgression between a pair of island endemic taxa, as well as ancient hybridization between their ancestors. Moreover, by comparing the relationship between introgression and recombination rate at these different time points, we found that selection against recent gene flow on the island is more prevalent than following ancient hybridization, likely due to the accumulation of reproductive barriers through time. Although it is possible that time has eroded previous signatures of selection against ancient introgression, we identified widespread phylogenetic discordance associated with ancient introgression, which suggests that much of the signal of ancient introgression has been preserved over time.

### 4.1 Recent and ancient hybridization contribute to distinct signals of introgression between a pair of island endemic monkeyflower subspecies

By examining the history of hybridization between these island taxa, we demonstrate how the diversity of taxa present within evolutionary radiations can be used to estimate the relative timing of past hybridization events. Martin et al. (2013) first demonstrated that it was possible to detect introgression that occurred further back in time by increasing the level of sequence divergence between the sister pair of ingroup taxa used in a four taxon test of introgression. Malinsky et al. (2018) advanced this approach by developing a statistic to estimate the relative timing of hybridization across a phylogenetic tree. In the current study, we further expand on these methods by performing two tests that can: 1) estimate levels of recent introgression in a genomic window, and 2) identify levels of ancient and recent introgression. These tests rely on the variation in the divergence times between a pair of hybridizing taxa and their respective sister taxa.

Specifically, we used Test 1 to demonstrate that hybridization between Island *longiflorus* and *parviflorus* began after Island and mainland *longiflorus* diverged from each other. This is because average levels of *f_d_* remain similar regardless of which taxon is used as P1 in the test (Fig 3B). We then performed additional tests for introgression using the sister pair of *aridus* and *parviflorus* as P1 and P2 to identify introgression that occurred prior to the divergence of clades C and D (Test 2). Because *parviflorus* and *aridus* are more diverged from each other than any of the taxa within clades C and D, Test 2 can detect recent hybridization between *parviflorus* and Island *longiflorus*, as well as any ancient introgression that occurred between the ancestor of *parviflorus* and the common ancestor of clades C and D. The difference between these two tests can be used to obtain an estimate of ancient introgression.

Although *parviflorus* is currently endemic to the Channel Islands, these results suggest that the ancestor of *parviflorus* was likely also present on the mainland of California, where it hybridized with the common ancestor of clades C and D at some point in the past. It has been argued that all plant species currently found on the Channel Islands are descended from mainland California ancestors (Axelrod 1965, Thorne 1969, Schoenherr et al. 2003). Furthermore, many of the woody plant species restricted to the Channel Islands were found on the mainland in the past, where they appear to have been driven to extinction by changing climate conditions (Axelrod 1965). Many of the Channel Island’s endemic species are believed to have migrated from the mainland during the Pleistocene glacial period when sea levels were lower (Johnson 1978, Muhs et al. 2015, Mychajliw et al. 2020). During this period, Santa Cruz Island was connected to Santa Rosa and Anacapa Islands, and the distance between these islands and the mainland was only between 6-10 km (Johnson 1978, Muhs et al. 2015), making dispersal to the islands possible. Furthermore, consistent with what we report here in *Mimulus*, there is also evidence of ancient and recent introgression between a pair of Channel Island endemic oak species (Ortego et al. 2018, Mead 2023, Mead et al. 2024). One of the island species appears to be a relic of a now extinct species that was present on the mainland, and the other is widely distributed on both the mainland and the Channel Islands. This raises the possibility that hybridization may be a common feature among closely related Channel Island endemics.

Indeed, hybridization between plant species on the Channel Islands appears to be common (Thorne 1969), which is consistent with a general finding of increased hybridization between closely related island endemics (Carlquist 1966, Reatini & Vision 2023). One proposed explanation for this is that hybridization can mitigate the effects of deleterious genetic load that accumulates in geographically isolated, small populations (Carlquist 1966). However, we observed no reductions in genetic diversity or effective population size in the island taxa, suggesting that there is no evidence of greater genetic load in either island taxon. Nevertheless, given that the samples of *parviflorus* and Island *longiflorus* sequenced here continue to display evidence of admixture, it is possible that these patterns of diversity and population size may be attributable to their shared history of introgression. Additional sampling from the other Channel Islands where the two taxa do not occur in sympatry will be necessary to test this hypothesis further.

### 4.2 Variation in the fitness effects of introgression over time

Speciation is a continuous process that involves the accumulation of reproductive isolation (Stankowski and Ravinet 2021). Multiple factors will determine the rate at which isolation evolves, but the time since divergence has been shown to be highly relevant (Coyne and Orr 1989). Thus, at any point along the speciation continuum before reproductive isolation is complete, hybrids can form, leading to the potential for gene flow between emerging species. The consequences of this gene flow for fitness will depend on the extent of isolation that has evolved and the environments that the taxa inhabit. For example, if a pair of taxa has only recently diverged, many of the variants exchanged between them will likely have few deleterious effects on fitness. However, as divergence times increase and more reproductive isolation becomes established, selection against hybrids may be stronger. Alternatively, variants transferred between taxa may be beneficial in a shared ecological environment. By identifying variation in the timing of introgression between taxa, we were able to reveal that the consequences of introgression for fitness also varied over time.

By building phylogenetic trees in windows across the genome, we found widespread phylogenetic discordance due to a combination of recent and ancient hybridization and incomplete lineage sorting. Surprisingly, we identified nearly equal numbers of windows that supported the species tree and the ancient introgression tree, suggesting that the signal of ancient introgression is widespread throughout the genome. In contrast, we identified a much smaller number of windows that supported the introgression tree, suggesting that recent introgression was limited to fewer regions throughout the genome.

To estimate the fitness effects of introgression, we compared the relationship between recombination rate and recent and ancient introgression. Selection against introgression should result in a positive relationship between introgression and local levels of recombination, because the recombination rate determines how quickly introgressed alleles will be separated from resident alleles. If selection acts against foreign variants, it should rapidly remove deleterious alleles in regions of low recombination where they remain linked with resident alleles (Brandvain et al. 2014, Schumer et al. 2018, Martin et al. 2019). Consistent with this expectation, we found that recent introgression between *parviflorus* and Island *longiflorus* was positively related to recombination rate, suggesting that admixed variants were often deleterious. By contrast, there was no relationship between ancient introgression and recombination rate. These findings imply that reproductive isolation has accumulated between *parviflorus* and Island *longiflorus*, such that more recent gene exchange between these diverged taxa resulted in selection against gene flow. By contrast, shortly after the split between their ancestors, there were likely fewer reproductive barriers in place, allowing free exchange of genetic information and a corresponding signal of neutral introgression. However, reproductive isolation is not complete between *parviflorus* and Island *longiflorus*, as numerous contemporary hybrids were detected in our samples (Fig 2B, 2C). Thus, additional work characterizing the components and extent of reproductive isolation, as well as the fitness of hybrids between these taxa will be needed to confirm these conclusions.

Another explanation for these findings is that the signatures of selection against ancient introgression have eroded over time. Although difficult to assess directly, we found widespread evidence of ancient introgression across the genome (Fig 4C), suggesting that much of this signal has been maintained through time. Indeed, the presence of numerous genomic windows with high support for the ancient introgression topology is consistent with anciently introgressed alleles being largely neutral. Although some windows likely display high support for the ancient introgression topology due to incomplete lineage sorting, windows of elevated “ancient *f_d_*” display strong support for the ancient introgression topology, suggesting that ancient introgression is at least partially contributing to this pattern. Moreover, many cases of widespread phylogenetic discordance (Nelson et al. 2021, Zhang et al. 2021a) and adaptive introgression (Meier, et al. 2017, Malinsky et al. 2018, Ma et al. 2019, Zhang et al. 2021b, Short & Streisfeld 2023) appear to be the result of ancient introgression, further implying that selection against introgression was weaker earlier in the divergence history of these taxa.

Finally, by examining *f_d_* values from Test 1 and Test 2 across the genome, we found extensive variation in genome wide levels of introgression, with some windows being nearly fixed for introgressed alleles. This raises the possibility that adaptation has contributed to the localized maintenance of introgressed alleles. Despite evidence that selection against recent gene flow was the primary factor shaping the heterogeneous patterns of introgression between *parviflorus* and Island *longiflorus*, we also found evidence for localized increases in the frequency of Q95 sites. Specifically, we identified a region on chromosome 5 in *parviflorus* that was nearly fixed for Island *longiflorus* alleles. Thus, although most alleles exchanged between these taxa likely decreased fitness, some introgressed alleles appear to have increased fitness. Both taxa inhabit similar environments and often occur in sympatry, implying that they experience many of the same selection pressures on Santa Cruz Island. Thus, the transfer of alleles from Island *longiflorus* into *parviflorus* may have been facilitated by their shared environments, but future studies on the ecology and physiology of these taxa will be needed to identify potential selective agents contributing to adaptation.

In conclusion, we identified evidence of selection against recent introgression between *parviflorus* and Island *longiflorus*, potentially because more reproductive barriers are currently in place. However, we found no current signal of selection between their ancestors. Thus, this study reveals that hybridization can occur at multiple points throughout the divergence history of a radiation, but the processes that shape their genomes can change over time.

## Data Accessibility Statement

Raw sequencing reads were downloaded from the Short-Read Archive (SRA) from bioproject ID: PRJNA549183. New sequencing reads generated here have been uploaded and added to bioproject ID PRJNA1149754. VCF files and population genomic data have been deposited to xxxx. The reference assembly and annotation are available at mimubase.org. Computer scripts used for population genomic analyses are available on Github at: https://github.com/awshort/Channel_Islands_monkeyflower_hybridization. Samples were collected from Santa Cruz Island according to the Scientific Research and Collecting Permit from the National Parks Service (CHIS-2024-SCI-0008).

## Benefit-Sharing Statement

Benefits generated: Benefits from this research accrue from the sharing of our data and results on public databases as described above.

## Author contributions

A.W.S. and M.A.S. designed the study, conducted all analyses, and wrote the manuscript.

## Acknowledgments

We would like to thank Jessie Crown for help in extracting the DNA for this project. We would also like to thank Peter Ralph, Andrew Kern, Yaniv Brandvain, and Bill Cresko for providing valuable feedback and discussion. We would also like to thank Doug Turnbull and Jason Carriere for preparing the libraries and conducting the Illumina sequencing at the UO Genomics & Cell Characterization Core Facility (GC3F). We would like to thank Dr. Cameron B. Williams for helping us to secure a scientific research and collection permit, and the National Park Service for granting us this permit.

## Supplemental Material

**Table S1.** Sampling locations and number of samples sequenced from each location for this study. Four additional samples from ECSC were sequenced in Stankowski et al. 2019.

| Population | number of samples | Latitude | Longitude |
| --- | --- | --- | --- |
| ECSC | 3 | 34.018 | -119.673 |
| SC2 | 8 | 34.01915 | -119.6804 |
| SC3 | 7 | 34.01925 | -119.68023 |
| SCB | 9 | 34.019267 | -119.68493 |

**Table S2.** The t-ratio from a linear mixed-effects model testing if the mean admixture proportion (*f_d_*) varied due to the divergence between the taxa used as P1 and P2 for the calculation of *f_d_*. The name of the taxon used as P1, and the mean taxonomic divergence (da) between the taxa used as P1 and P2 are presented. Statistical significance is denoted as: *** for p < 0.001, ** for p < 0.01, and * for p < 0.05.

| Factor Level | Estimate | df | t-ratio (Fd) |
| --- | --- | --- | --- |
| *aridus* (0.00868) vs *aurantiacus* (0.00409) | 0.02629 | 10587 | 4.621*** |
| *aridus* (0.00868) vs *calycinus* (0.00236) | 0.03681 | 10587 | 6.588*** |
| *aridus* (0.00868) vs *longiflorus* (0.00219) | 0.035 | 10587 | 6.269*** |
| *aridus* (0.00868) vs OC (0.00333) | 0.03209 | 10587 | 5.726*** |
| *aridus* (0.00868) vs red (0.00328) | 0.032 | 10587 | 5.709*** |
| *aridus* (0.00868) vs yellow (0.00311) | 0.03233 | 10587 | 5.765*** |
| *aurantiacus* (0.00409) vs *calycinus* (0.00236) | 0.01052 | 10587 | 1.831 |
| *aurantiacus* (0.00409) vs *longiflorus* (0.00219) | 0.00871 | 10587 | 1.517 |
| *aurantiacus* (0.00409) vs OC (0.00333) | 0.0058 | 10587 | 1.007 |
| *aurantiacus* (0.00409) vs red (0.00328) | 0.00571 | 10587 | 0.991 |
| *aurantiacus* (0.00409) vs yellow (0.00311) | 0.006 | 10587 | 1.048 |
| *calycinus* (0.00236) vs *longiflorus* (0.00219) | -0.00181 | 10587 | -0.32 |
| *calycinus* (0.00236) vs OC (0.00333) | -0.00472 | 10587 | -0.833 |
| *calycinus* (0.00236) vs red (0.00328) | -0.00481 | 10587 | -0.849 |
| *calycinus* (0.00236) vs yellow (0.00311) | -0.00448 | 10587 | -0.79 |
| *longiflorus* (0.00219) vs OC (0.00333) | -0.00291 | 10587 | -0.514 |
| *longiflorus* (0.00219) vs red (0.00328) | -0.003 | 10587 | -0.53 |
| *longiflorus* (0.00219) vs yellow (0.00311) | -0.00267 | 10587 | -0.471 |
| OC (0.00333) vs red (0.00328) | -0.00009 | 10587 | -0.016 |
| OC (0.00333) vs yellow (0.00311) | 0.00024 | 10587 | 0.042 |
| red (0.00328) vs yellow (0.00311) | 0.00033 | 10587 | 0.058 |

**Table S3.** The t-ratio from a linear mixed-effects model testing if admixture proportion (*f_d_*) varied due to the divergence between Island *longiflorus* and the taxon used as P3 for the calculation of *f_d_*. The name of the taxon used as P3 and the mean taxonomic divergence between Island *longiflorus* and the taxon used as P3 for the calculation of *f_d_* are presented. Statistical significance is denoted as: *** for p < 0.001, ** for p < 0.01, and * for p < 0.05.

| Factor Level | Estimate | df | t-ratio (Fd) |
| --- | --- | --- | --- |
| *aurantiacus* (0.00409) vs *calycinus* (0.00236) | -0.01163 | 9044 | -2.54 |
| *aurantiacus* (0.00409) vs CI *longiflorus* (0.0) | -0.06997 | 9044 | -16.035*** |
| *aurantiacus* (0.00409) vs *longiflorus* (0.00219) | -0.01129 | 9044 | -2.48 |
| *aurantiacus* (0.00409) vs OC (0.00333) | -0.00909 | 9044 | -1.942 |
| *aurantiacus* (0.00409) vs red (0.00328) | -0.01068 | 9044 | -2.248 |
| *aurantaicus* (0.00409) vs yellow (0.00311) | -0.01018 | 9044 | -2.13 |
| *calycinus* (0.00236) vs CI *longiflorus* (0.0) | -0.05833 | 9044 | -13.188*** |
| *calycinus* (0.00236) vs *longiflorus* (0.00219) | 0.00034 | 9044 | 0.074 |
| *calycinus* (0.00236) vs OC (0.00333) | 0.00254 | 9044 | 0.536 |
| *calycinus* (0.00236) vs red (0.00328) | 0.00096 | 9044 | 0.199 |
| *calycinus* (0.00236) vs yellow (0.00311) | 0.00145 | 9044 | 0.301 |
| CI *longiflorus* (0.0) vs *longiflorus* (0.00219) | 0.05867 | 9044 | 13.351*** |
| CI *longiflorus* (0.0) vs OC (0.00333) | 0.06087 | 9044 | 13.443*** |
| CI *longiflorus* (0.0) vs red (0.00328) | 0.05929 | 9044 | 12.897*** |
| CI *longiflorus* (0.0) vs yellow (0.00311) | 0.05978 | 9044 | 12.917*** |
| *longiflorus* (0.00219) vs OC (0.00333) | 0.0022 | 9044 | 0.467 |
| *longiflorus* (0.00219) vs red (0.00328) | 0.00062 | 9044 | 0.129 |
| *longiflorus* (0.00219) vs yellow (0.00311) | 0.00111 | 9044 | 0.231 |
| OC (0.00333) vs red (0.00328) | -0.00158 | 9044 | -0.323 |
| OC (0.00333) vs yellow (0.00311) | -0.00109 | 9044 | -0.221 |
| red (0.00328) vs yellow (0.00311) | 0.005 | 9044 | 0.099 |

**Table S4.** The t-ratio from a linear mixed-effects model testing if the mean admixture proportion for Test 1 *f_d_* varies among recombination rate quantile bins. Quantile bins of recombination rate (in cM/Mb) are presented. Statistical significance is denoted as: *** for p < 0.001, ** for p < 0.01, and * for p < 0.05. Calculations of Test 1 *f_d_* were performed using mainland *longiflorus* as P1, Island *longiflorus* as P2, and *parviflorus* as P3. *M. clevelandii* was used as the outgroup.

| Factor Level | Estimate | df | t-ratio (Fd) |
| --- | --- | --- | --- |
| [0,0.824] vs (0.824,1.51] | 0.04604 | 1463 | 3.419** |
| [0,0.824] vs (1.51,2.33] | 0.04239 | 1463 | 3.156* |
| [0,0.824] vs (2.33,3.66] | 0.04099 | 1463 | 3.085* |
| [0,0.824] vs (3.66,10.9] | 0.07756 | 1463 | 5.939*** |
| (0.824,1.51] vs (1.51,2.33] | 0.00365 | 1463 | 0.271 |
| (0.824,1.51] vs (2.33,3.66] | 0.00505 | 1463 | 0.380 |
| (0.824,1.51] vs (3.66,10.9] | -0.03152 | 1463 | -2.411 |
| (1.51,2.33] vs (2.33,3.66] | 0.00140 | 1463 | 0.106 |
| (1.51,2.33] vs (3.66,10.9] | -0.03517 | 1463 | -2.698 |
| (2.33,3.66] vs (3.66,10.9] | -0.03657 | 1463 | -2.838* |

**Table S5.** The t-ratio from a linear mixed-effects model testing if the Test 2 admixture proportion (*fd*) varies among recombination rate quantile bins. Quantile bins of recombination rate (in cM/Mb) are presented. Statistical significance is denoted as: *** for p < 0.001, ** for p < 0.01, and * for p < 0.05. Calculations of Test 2 *f_d_* were performed using *aridus* as P1, *parviflorus* as P2, and Island *longiflorus* as P3. *M. clevelandii* was used as the outgroup.

| Factor Level | Estimate | df | t-ratio (Fd) |
| --- | --- | --- | --- |
| [0,0.824] vs (0.824,1.51] | 0.02829 | 1514 | 2.322 |
| [0,0.824] vs (1.51,2.33] | 0.02597 | 1514 | 2.114 |
| [0,0.824] vs (2.33,3.66] | 0.02086 | 1514 | 1.739 |
| [0,0.824] vs (3.66,10.9] | 0.05104 | 1514 | 4.258*** |
| (0.824,1.51] vs (1.51,2.33] | 0.00232 | 1514 | 0.191 |
| (0.824,1.51] vs (2.33,3.66] | 0.00744 | 1514 | 0.626 |
| (0.824,1.51] vs (3.66,10.9] | -0.02275 | 1514 | -1.918 |
| (1.51,2.33] vs (2.33,3.66] | 0.00511 | 1514 | 0.427 |
| (1.51,2.33] vs (3.66,10.9] | -0.02507 | 1514 | -2.095 |
| (2.33,3.66] vs (3.66,10.9] | -0.03018 | 1514 | -2.587 |

**Table S6.** The t-ratio from a linear mixed-effects model testing if the ancient admixture proportion (*f_d_*) varies among recombination rate quantile bins. Quantile bins of recombination rate (in cM/Mb) are presented. Statistical significance is denoted as: *** for p < 0.001, ** for p < 0.01, and * for p < 0.05. Ancient *f_d_* was calculated by taking the difference between *f_d_* calculated from Test 2 and Test 1.

| Factor Level | Estimate | df | t-ratio (Fd) |
| --- | --- | --- | --- |
| [0,0.824] vs (0.824,1.51] | -0.00643 | 1049 | -0.633 |
| [0,0.824] vs (1.51,2.33] | -0.01527 | 1049 | -1.506 |
| [0,0.824] vs (2.33,3.66] | -0.01299 | 1049 | -1.303 |
| [0,0.824] vs (3.66,10.9] | -0.01479 | 1049 | -1.433 |
| (0.824,1.51] vs (1.51,2.33] | 0.00884 | 1049 | 0.862 |
| (0.824,1.51] vs (2.33,3.66] | 0.00656 | 1049 | 0.650 |
| (0.824,1.51] vs (3.66,10.9] | 0.00836 | 1049 | 0.801 |
| (1.51,2.33] vs (2.33,3.66] | -0.00229 | 1049 | -0.227 |
| (1.51,2.33] vs (3.66,10.9] | -0.00048 | 1049 | -0.046 |
| (2.33,3.66] vs (3.66,10.9] | 0.00181 | 1049 | 0.176 |

**Figure S1.**
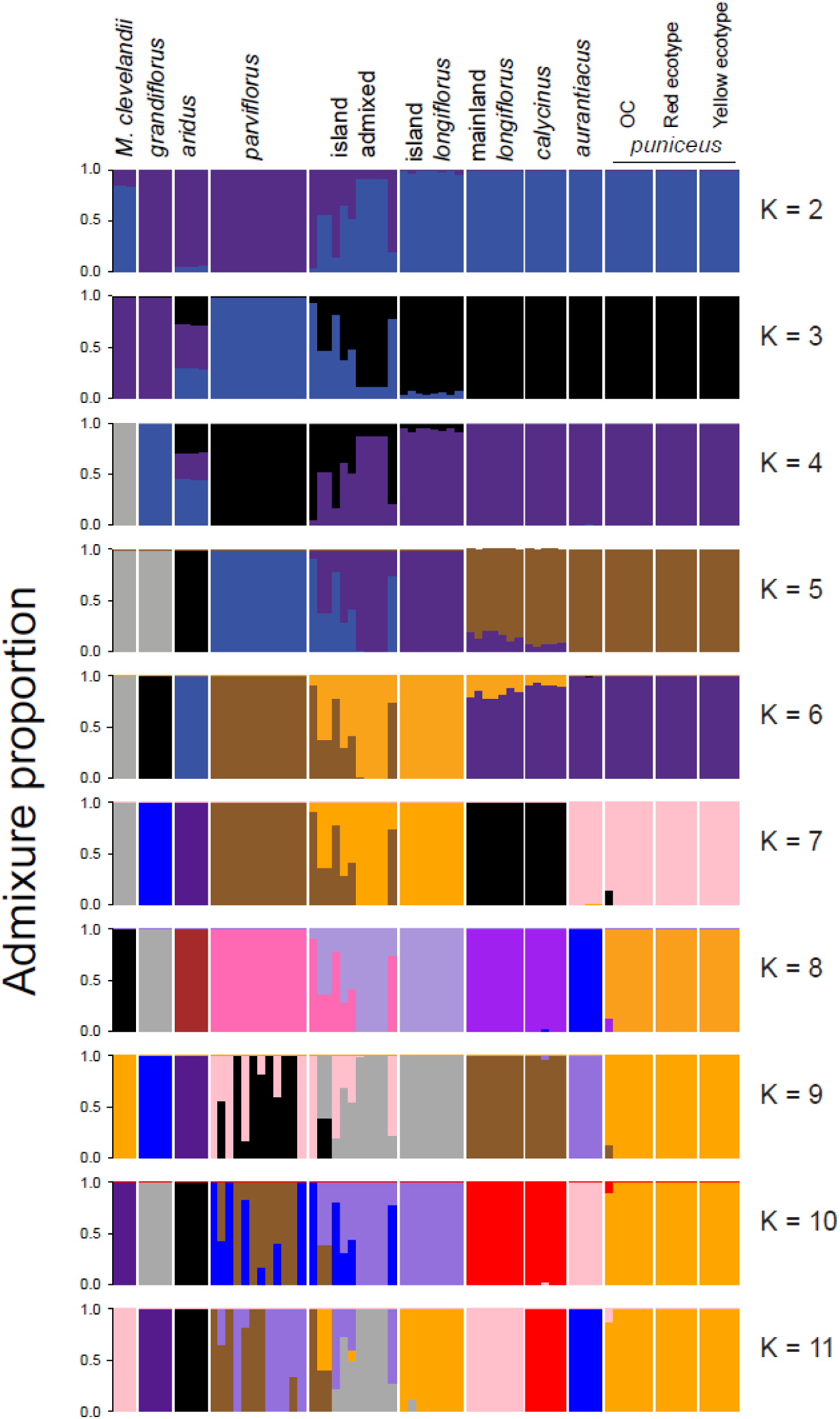
The ancestry proportions from *Admixture* at K = 2 to K=11 for samples from all the subspecies and their sister species, *M. clevelandii*.

**Figure S2.**
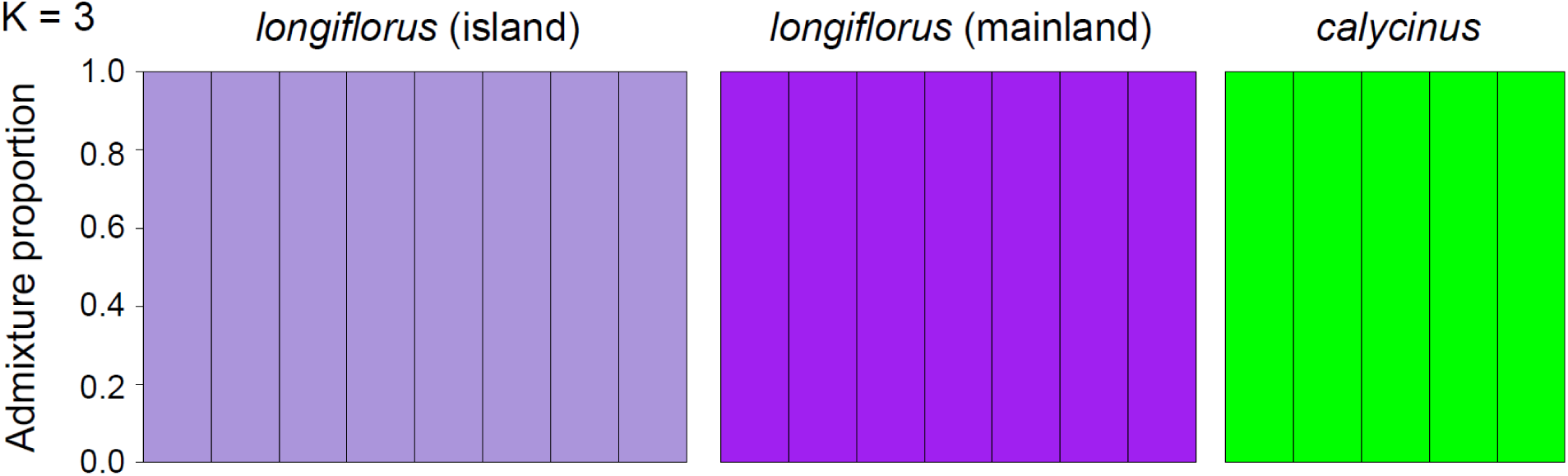
The ancestry proportions from *Admixture* at K = 3 from a run that included only the island *longiflorus*, mainland *longiflorus*, and *calycinus* samples.

**Figure S3.**
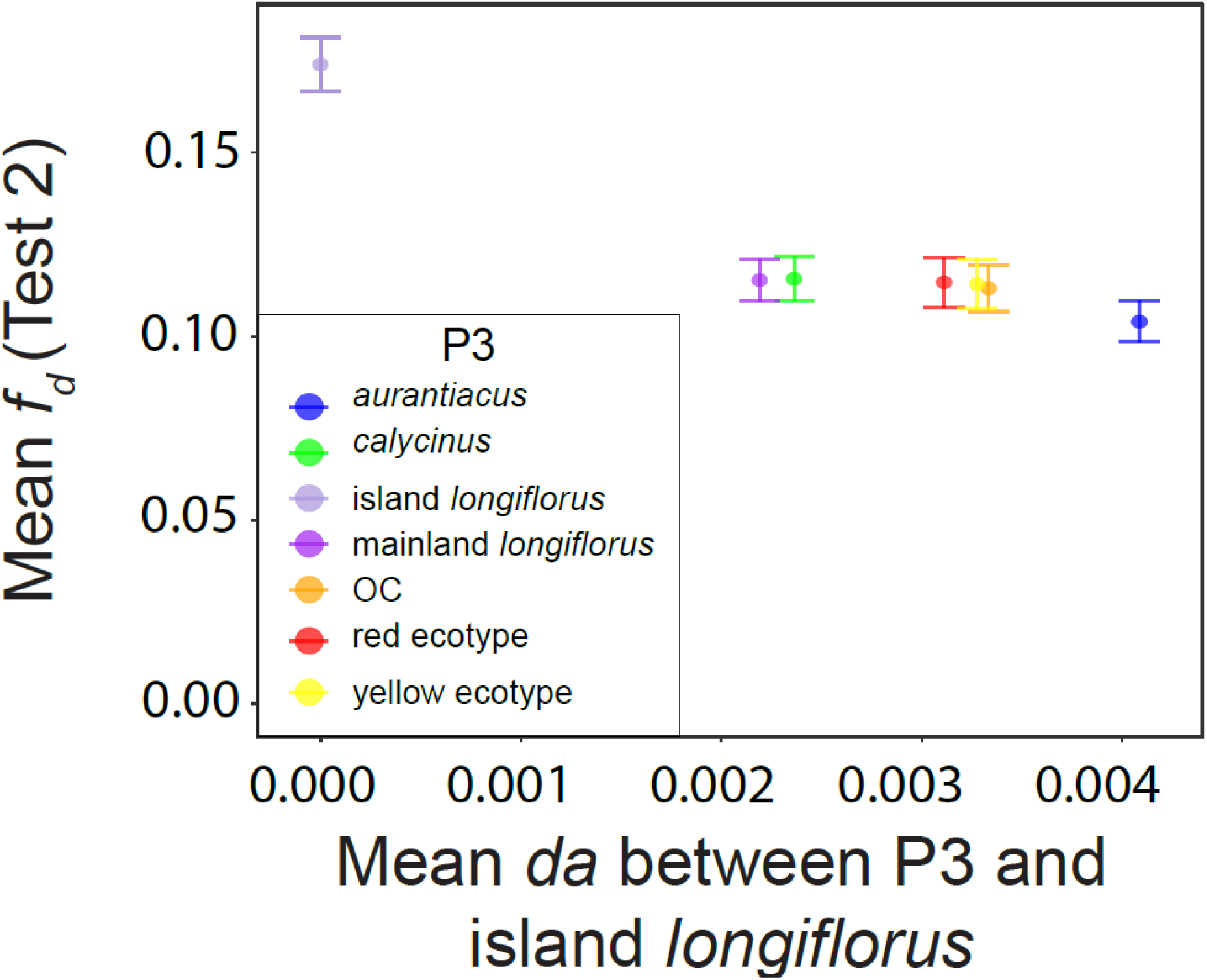
Mean and 95% confidence intervals of Test 2 *f_d_* values calculated in 100kb windows are plotted against mean levels of sequence divergence (da) between Island *longiflorus* and the clade C or D taxa used as P3 for the calculation of *f_d_*. Test 2 calculations of fd were performed using *aridus* as P1, *parviflorus* as P2, and the various clade C and D taxa as P3. Colors indicate the P3 taxon used to calculate *f_d_*.

**Figure S4.**
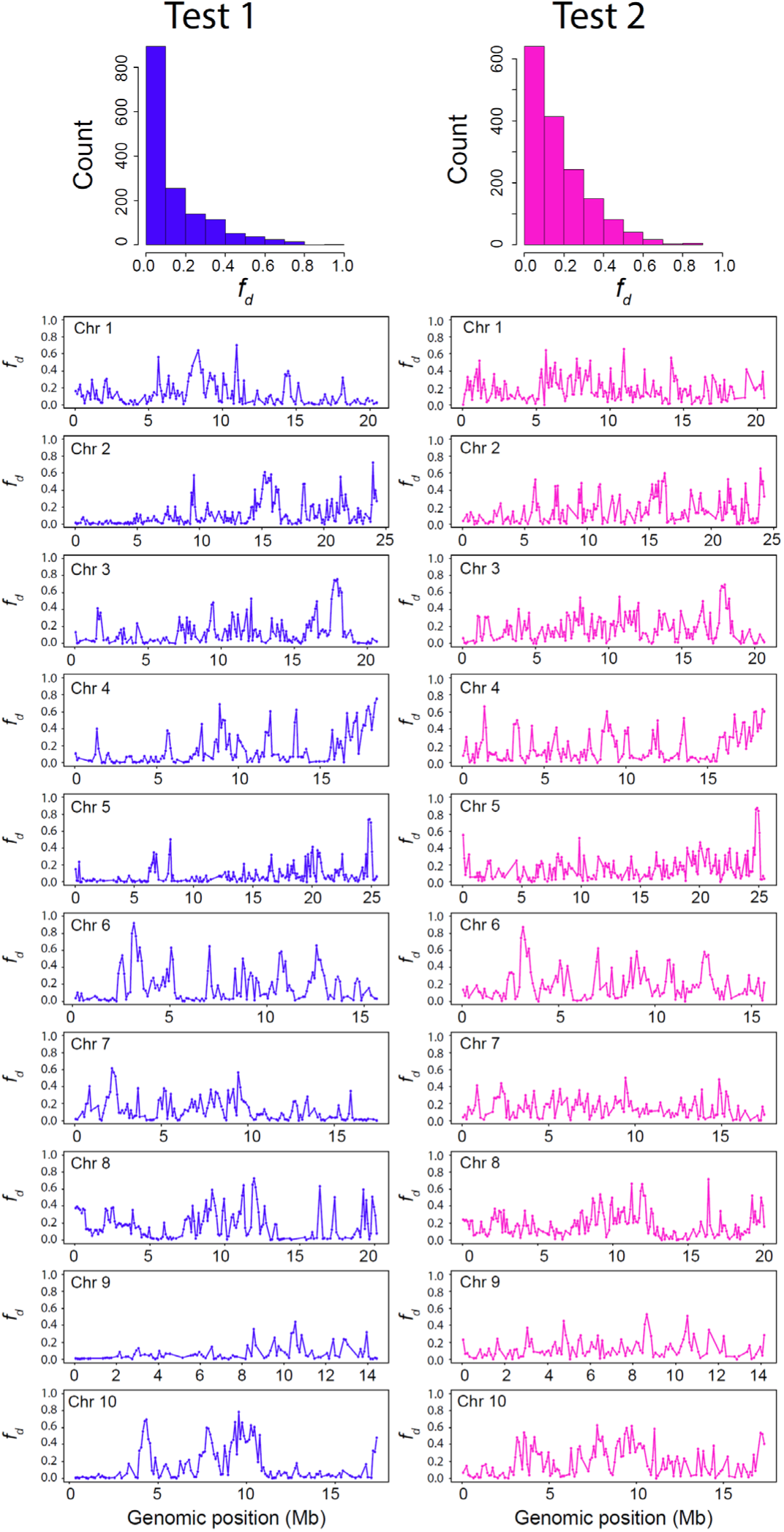
Histogram of the distribution of the raw Test 1 (blue) and Test 2 (pink) *f_d_* values calculated in 100 kb windows. Genome wide variation of raw *f_d_* values plotted across the 10 chromosomes of the *M. aurantiacus* genome. Test 1 *f_d_* values are in blue and Test 2 *f_d_* values are in pink. Test 1 calculations of *f_d_* were performed using mainland *longiflorus* as P1, island *longiflorus* as P2, and *parviflorus* as P3. Test 2 calculations of *f_d_* were performed using *aridus* as P1, *parviflorus* as P2, and island *longiflorus* as P3.

**Figure S5.**
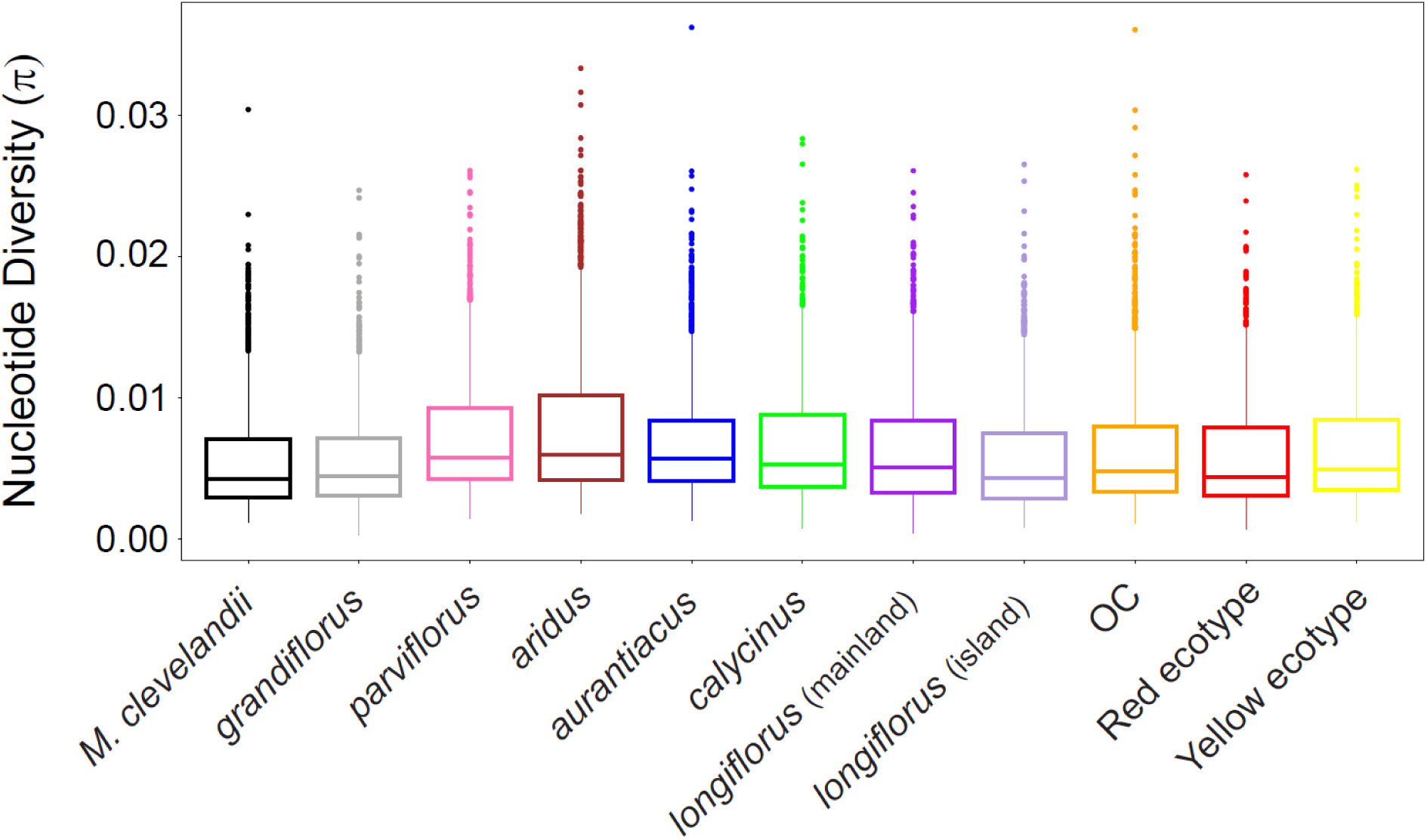
Boxplots of the distribution of nucleotide diversity (π) calculated in 100 kb windows for all the subspecies and their sister species, *M. clevelandii*.

**Figure S6.**
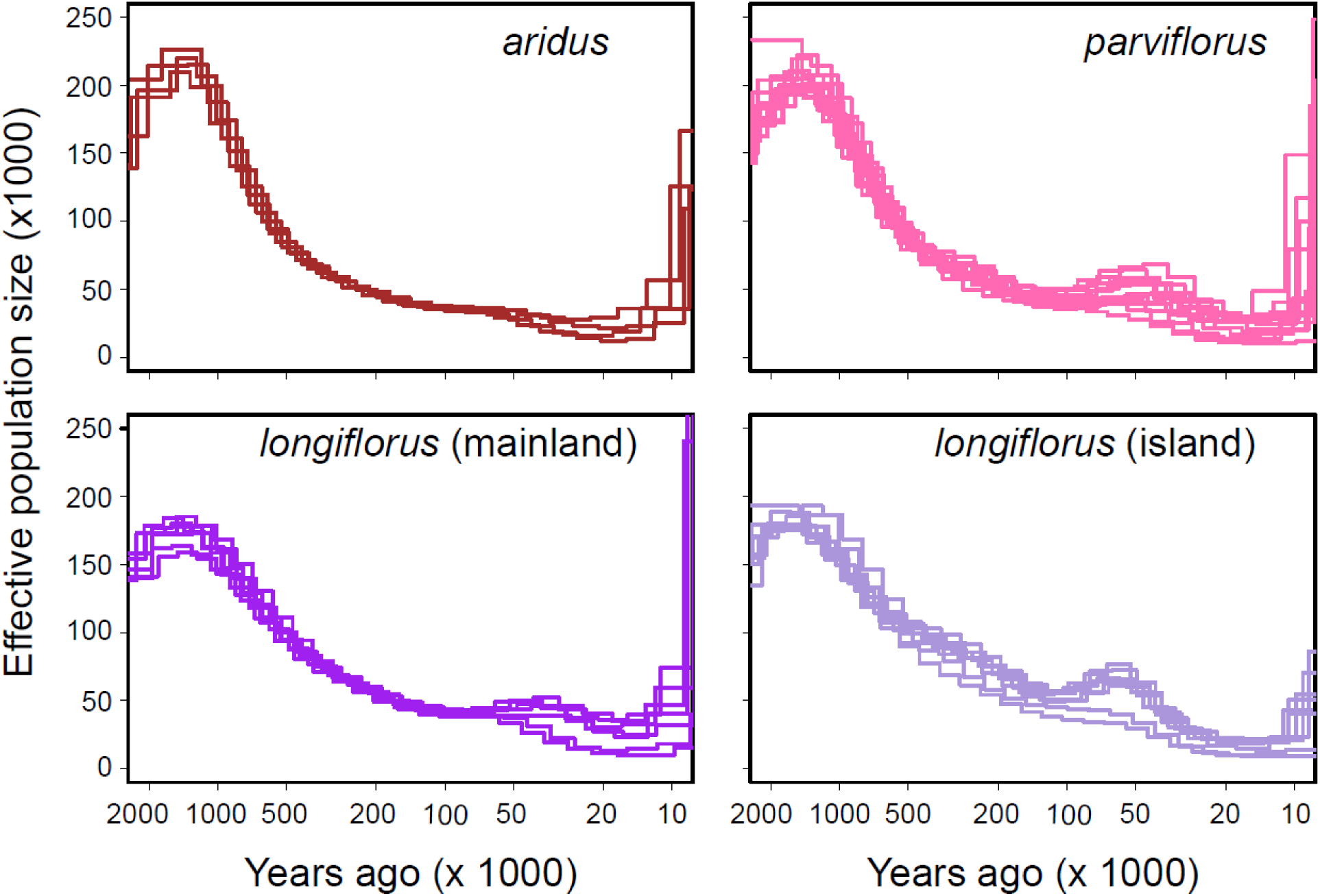
The estimated variation in effective population size over time from PSMC for each sequenced sample of *aridus*, *parviflorus*, mainland *longiflorus*, and island *longiflorus*.

**Figure S7.**
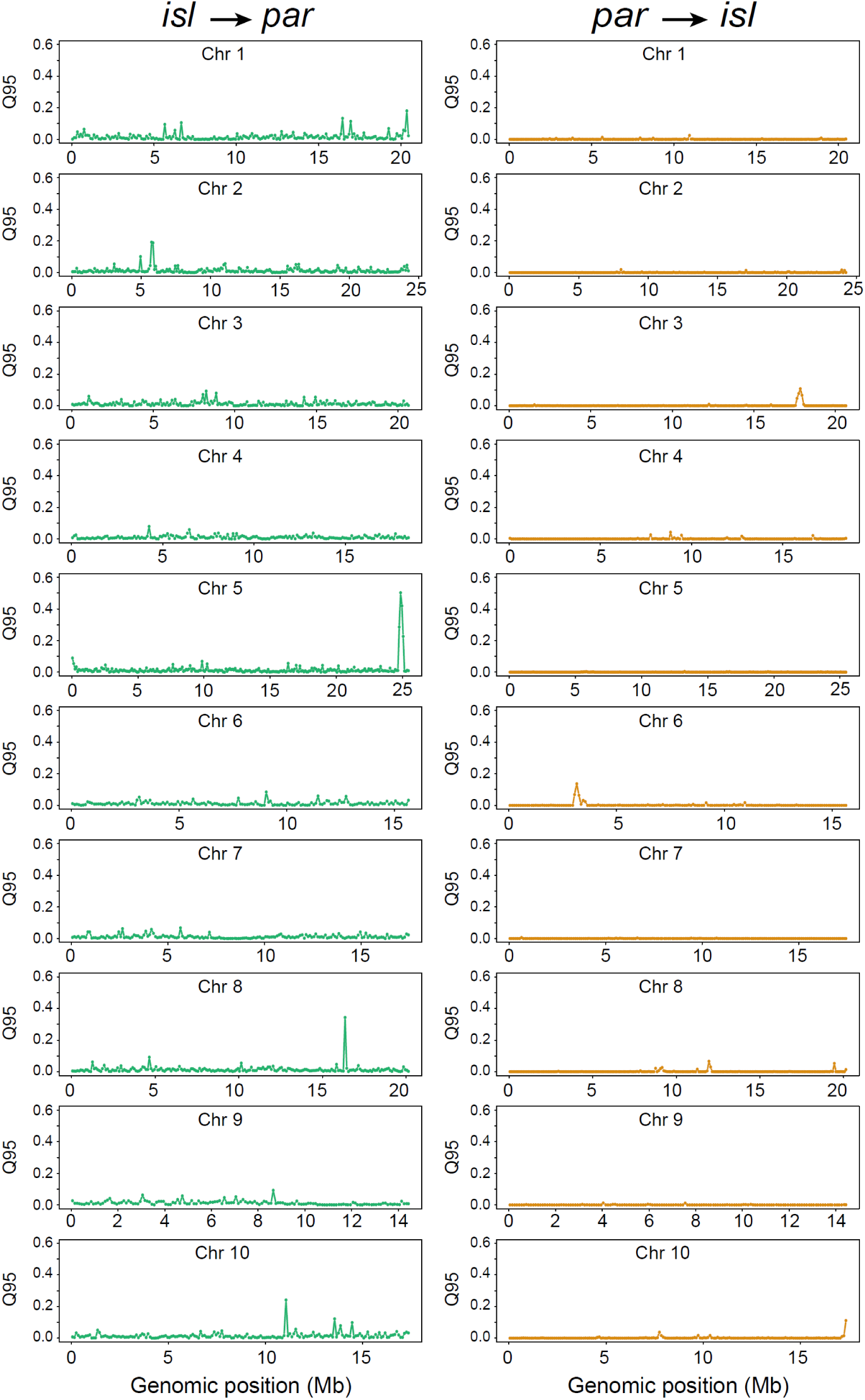
Genome wide variation in the Q95 statistic calculated in 100 kb windows plotted across the 10 chromosomes of the *M. aurantiacus* genome. Colors indicate the directionality of introgression, with gene flow from Island *longiflorus* into *parviflorus* in green, and gene flow from *parviflorus* into Island *longiflorus* in yellow.

**Figure S8.**
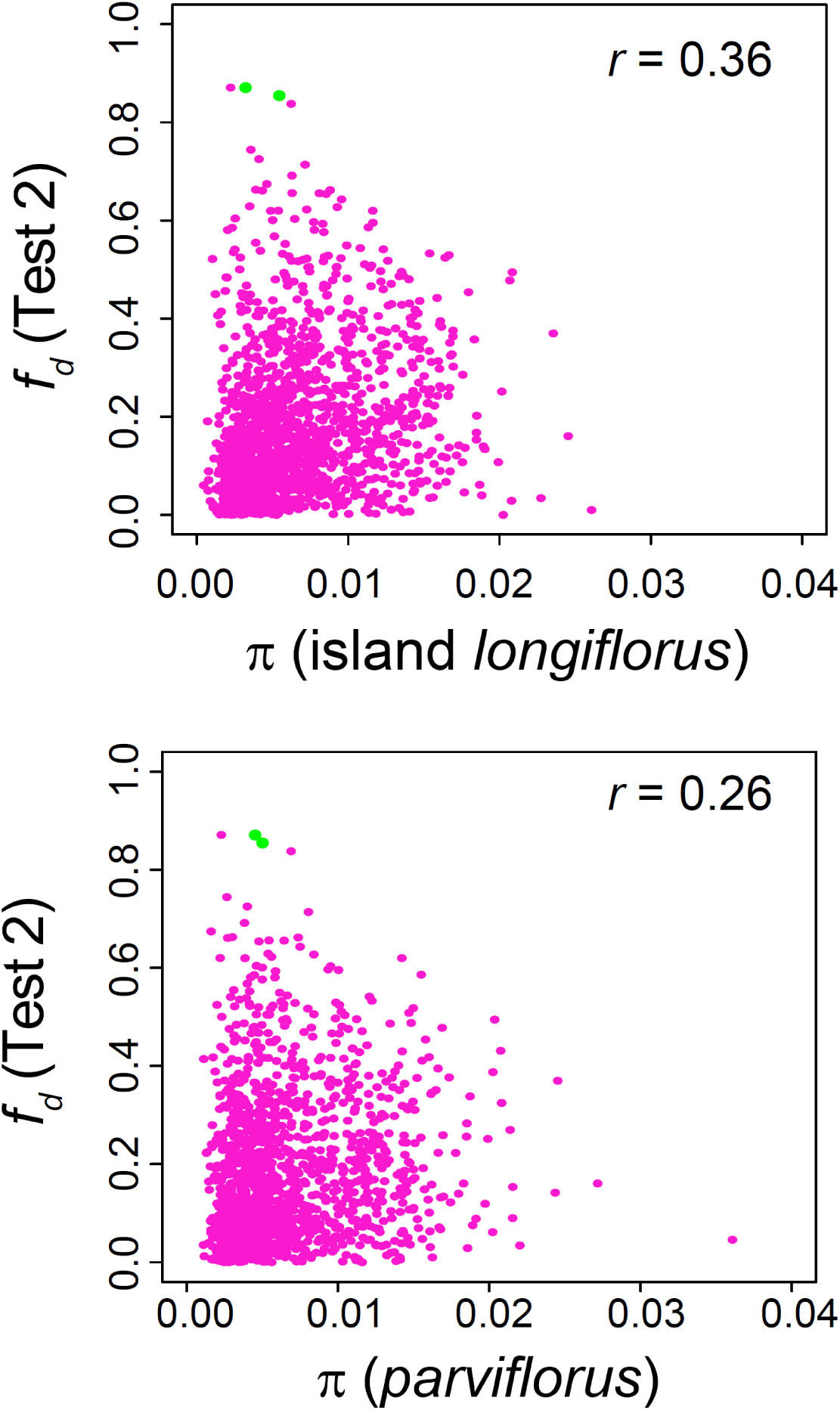
Scatterplots showing the relationship between Test 2 *f_d_* and nucleotide diversity (π) for Island *longiflorus* (top) and *parviflorus* (bottom). The correlation coefficient between the statistics is presented in the upper right-hand corner of each plot. Test 2 calculations of fd were performed using *aridus* as P1, *parviflorus* as P2, and Island *longiflorus* as P3. Green points indicate the windows with the highest *f_d_* and Q95 on chromosome 5.

